# RNase L-mediated RNA decay alters 3’ end formation and splicing of host mRNAs

**DOI:** 10.1101/2022.01.28.478180

**Authors:** James M. Burke, Nina Ripin, Max B. Ferretti, Laura A. St Clair, Emma R. Worden-Sapper, Sara L. Sawyer, Rushika Perera, Kristen W. Lynch, Roy Parker

## Abstract

The antiviral endoribonuclease, RNase L, is a vital component of the mammalian innate immune response that destroys host and viral RNA to reduce viral gene expression. Herein, we show that a consequence of RNase L-mediated decay of cytoplasmic host RNAs is the widespread re-localization of RNA-binding proteins (RBPs) from the cytoplasm to the nucleus, due to the presence of nuclear RNA. Concurrently, we observe global alterations to host RNA processing in the nucleus, including alterations of splicing and 3’ end formation, with the latter leading to downstream of gene (DoG) transcripts. While affecting many host mRNAs, these alterations are pronounced in mRNAs encoding type I and type III interferons and coincide with the retention of their mRNAs in the nucleus. Similar RNA processing defects also occur during infection with either dengue virus or SARS-CoV-2 when RNase L is activated. These findings reveal that the distribution of RBPs between the nucleus and cytosol is fundamentally dictated by the availability of RNA in each compartment and thus viral infections that trigger cytoplasmic RNA degradation alter RNA processing due to the nuclear influx of RNA binding proteins.

## INTRODUCTION

Ribonuclease L (RNase L) limits the replication of diverse viruses, including influenza virus, chikungunya virus, SARS-CoV-2, and vaccinia virus (Min and Krug, 2006; Cooper et al., 2015; Li et al., 2016; Li et al., 2021). RNase L is activated by 2’-5’-oligo(A), which is produced by 2’-5’-oligoadenylate synthetases (OASs) upon binding to viral or endogenous dsRNA (reviewed in Silverman, 2007; Kristiansen et al., 2011). RNase L cleaves at UN^N motifs in single-stranded RNA (ssRNA) regions (Floyd-Smith et al., 1981; Wreschner et al., 1981), and destruction of viral RNAs by RNase L reduces viral replication (Han et al., 2007; Cooper et al., 2015, Nogimori et al., 2019; Burke et al., 2021b.). In addition, RNase L also cleaves several types of cellular RNAs, including rRNAs, tRNAs, and mRNAs (Wreschner et al., 1981; Andersen et al, 2009; Donovan et al., 2017; Rath et al., 2019; Burke et al, 2019).

We, and others, recently demonstrated that RNase L activation results in widespread degradation of most host basal mRNAs (Burke et al., 2019; Rath et al., 2019). This has several impacts to cellular RNA biology. First, it allows for antiviral programming of translation since several immediate early antiviral mRNAs (i.e., *interferon* mRNAs) escape RNase L-mediated mRNA decay. Second, it inhibits the assembly of stress granules (Burke et al., 2020), cytoplasmic ribonucleoprotein (RNP) complexes proposed to modulate aspects of the antiviral response (Onomoto et al., 2012; Reineke and Lloyd, 2014; Yoo et al., 2014). Third, it promotes the formation of an alternative cytoplasmic RNP granule of unknown function termed RNase L-dependent bodies (RLBs) (Burke et al., 2020). Fourth, it inhibits nuclear mRNA export, which substantially reduces protein production of influenza virus as well as host cytokines induced by dsRNA, dengue virus infection, or SARS-CoV-2 infection (Burke et al., 2021a, Burke et al., 2021b). Lastly, it results in the translocation of poly(A)-binding protein (PABP) from the cytoplasm to the nucleus (Burke et al., 2019).

Herein, we demonstrate that RNase L-mediated degradation of host mRNAs primarily occurs in the cytoplasm, which leads to the re-localization of many RBPs to the nucleus in a manner dependent on intact nuclear RNA. This demonstrates the fundamental principle that the distribution of RBPs between subcellular compartments is dependent on the availability of their binding sites. The influx of RBPs into the nucleus is concurrent with global RNase L-dependent RNA processing alterations including alternative splicing, intron retention and transcription read-through leading to downstream of gene (DoG) transcript production, both of which correlate with inhibition of nuclear export of *IFN* mRNAs. These are general responses since we observe these alterations in RNA processing occur during exposure to exogenous dsRNA, dengue virus serotype 2 (DENV2) infection, or severe acute respiratory syndrome coronavirus 2 (SARS-CoV-2) infection. These findings show that RNase L-mediated mRNA decay alters the balance of RNA-binding protein subcellular localization, host RNA processing events, and antiviral gene expression. Since RBPs can also regulate transcription (Xiao et al., 2019), this work strongly implies that widespread RNA degradation in the cytosol will also lead to changes in transcription of multiple genes.

## RESULTS

### RNase L activation triggers re-localization of multiple RNA-binding proteins to the nucleus

The poly(A)-binding protein (PABP) primarily localizes to the cytoplasm in unstressed A549 cells (Figure 1A). However, upon lipofection of poly(I:C), PABP translocates to the nucleus (Fig. 1A & Burke et al., 2019). The translocation of PABP to the nucleus in response to poly(I:C) is dependent on RNase L since it does not occur in RNase L-KO A549 cells (Fig. 1A) but is rescued in RNase L-KO cells upon restoration of RNase L expression (Burke et al., 2019).

**Fig. 1.**
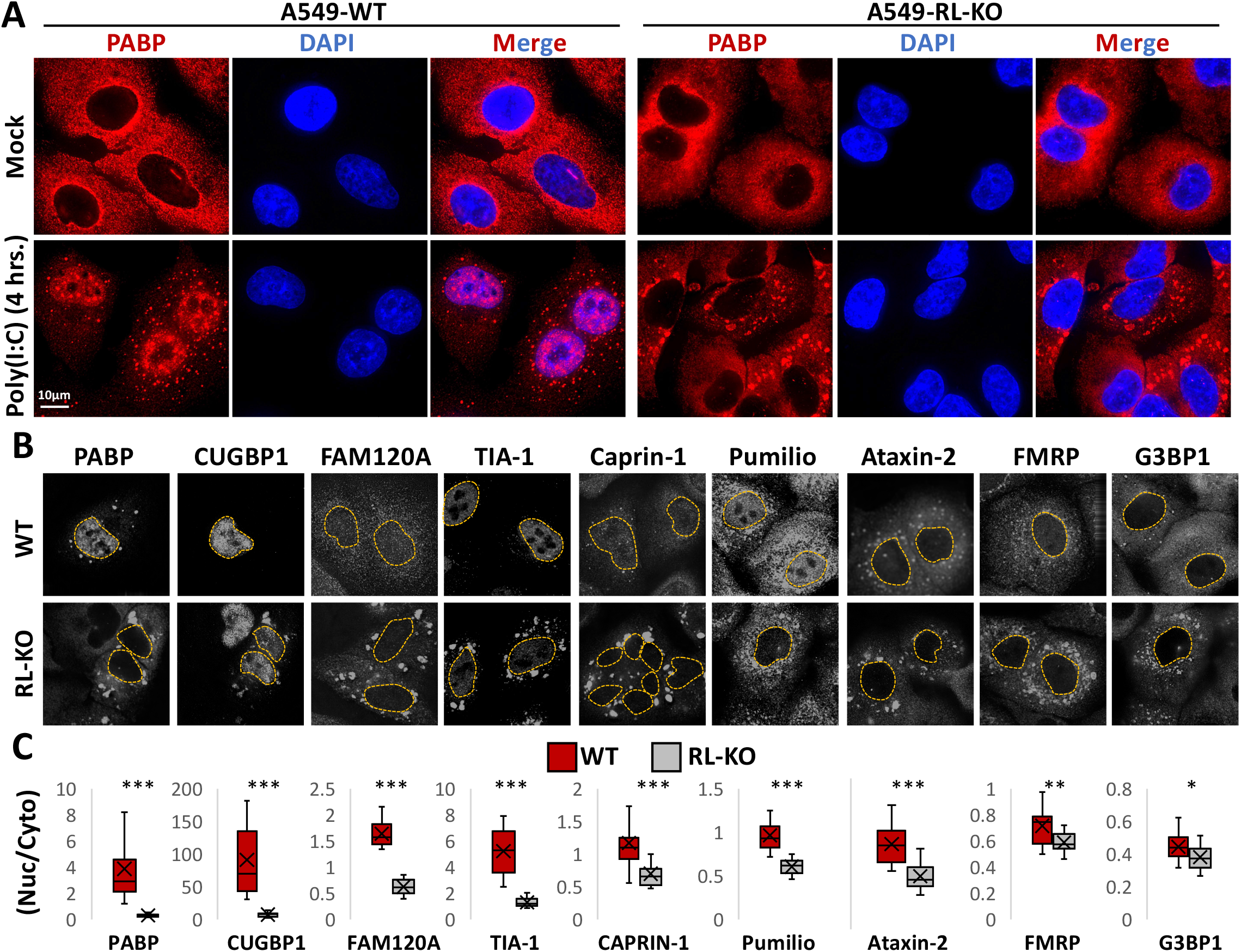
RNase L-dependent accumulation of RBPs in the nucleus. (A) Immunofluorescence assay for PABP in wild-type (WT) or RNase L knockout (RL-KO) A549 cells following four hours of lipofection with or without poly(I:C). (B) Immunofluorescence assay for RNA-binding proteins that enrich in cytoplasmic stress granules. Dashed line represents the nuclear boundary as determined by DAPI staining. (C) Quantification of the ratio (nucleus/cytoplasm) of the mean fluorescence intensity.

To determine if RNase L activation led to the import of additional RNA-binding proteins (RBPs), we performed immunofluorescence assays for several RBPs in WT and RL-KO cells post-poly(I:C) lipofection. A striking result is that RNase L activation promotes the nuclear localization of several RBPs (Fig. 1B,C), including CUGBP-1, FAM120A, TIA-1, Caprin-1, pumilio, and Ataxin-2, although some RBPs, such as G3BP1 and FMRP, only showed small, but significant, increases (Fig. 1B,C). These data indicate that RNase L activation results in the accumulation of RBPs in the nucleus.

#### RNase L and RNase L-mediated RNA decay are primarily localized to the cytoplasm

One mechanism by which RNase L could alter the localization of RBPs between the nucleus and cytoplasm is differential RNA degradation between the cytoplasm and nucleus, whereby higher RNA decay in the cytoplasm relative to the nucleus would lead to disassociation of RBPs from cytoplasmic RNA more rapidly than from nuclear RNA. To assess this, we quantified poly(A)+ RNA in the cytoplasm and nucleus of WT and RL-KO cells lipofected with or without post-poly(I:C) (Fig. 2A). We also stained cells for G3BP1 to identify cells undergoing a dsRNA response, whereby WT cells with activated RNase L contain RNase L-dependent bodies (RLBs) or RL-KO cells with activated PKR contain stress granules (SGs) (Burke et al., 2020).

**Fig. 2.**
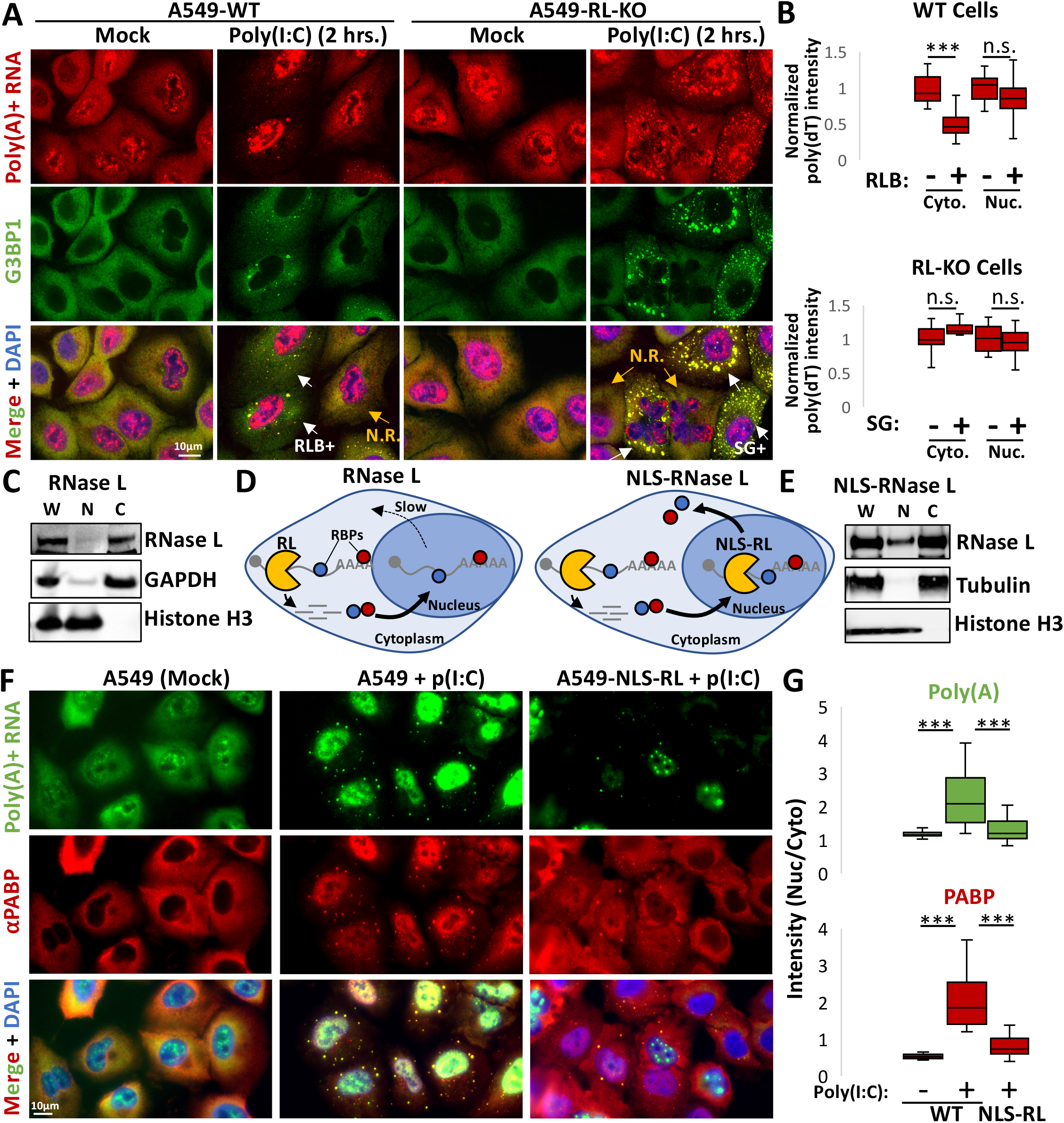
Decay of RNAs specifically in the cytoplasm by RNase L results in nuclear accumulation of RBPs. (A) FISH for poly(A)+ RNA and immunofluorescence assay for G3BP1 in WT and R-KO A549 cells. WT cells with RNase L-dependent bodies (RLBs) or RL-KO cells containing stress granules (SGs) are indicated by white arrows. Cells without RLBs or SGs are non-responsive (N.R.) and are indicated by yellow arrows. (B) Quantification of poly(A)+ RNA signal in the nucleus (nuc.) or cytoplasm (cyto.) in WT and RL-KO cells with or without RLBs or SGs, respectively, as represented in (A). (C) Immunoblot analysis of endogenous RNase L in whole cell lysate (w), nuclear fraction (n), or cytoplasmic fraction (c) from A549-WT cells. (D) Schematic showing RBP re-localization following either cytoplasmic RNase L activation (left) or activation of nuclear-localized RNase L (RL-NLS). (E) Immunoblot for RNase L in whole cell (w), nuclear (n), and cytoplasmic (c) crude fractions showing nuclear localization of the RNase L-NLS. (F) Immunofluorescence assay for PABP and FISH for poly(A)+ RNA in parental A549 cells or A549 that express RNase L-NLS construct following mock or poly(I:C) lipofection. (G) Quantification of PABP and poly(A)+ RNA signal from (F).

Importantly, we did not observe a significant reduction of poly(A)+ RNA staining in the nucleus of RLB-positive WT cells (Fig. 2A,B), whereas poly(A)+ RNA was significantly reduced in the cytoplasm (Fig. 2A,B). Moreover, we did not observe reduced cytoplasmic poly(A)+ RNA staining in SG-positive RL-KO cells. Consistent with these observations, the *GAPDH* mRNA accumulates in the nucleus while being degraded in the cytoplasm when RNase L is activated (Burke et al., 2021a). These data indicate that RNase L-mediated RNA decay primarily occurs in the cytoplasm in A549 cells.

While previous studies have shown that RNase L can localize to both the cytoplasm and nucleus (Bayard and Gabrion, 1993; Al-Ahmadi et al., 2009), we observed that endogenous RNase L is almost exclusively localized to the cytoplasm in A549 cells via cellular fractionation followed by immunoblot analysis (Fig. 2C). Combined, these data indicate that RNase L-mediated RNA decay primarily occurs in the cytoplasm.

#### Nuclear influx of RBPs is dependent on nuclear RNA

In principle, RNase L activation could alter RBP localization in two manners. First, RBPs may be subject to post-translational modifications, indirectly promoted by RNase L activation, that increase their nuclear accumulation. Alternatively, free RBPs may shuttle between the nucleus and cytosol faster that RBPs bound to RNA, and therefore the degradation of bulk cytoplasmic RNA would allow RBPs to shuttle to the nucleus, where binding to nuclear RNA would retain the RBPs in the nucleus. A prediction of this latter model is that the accumulation of nuclear RBPs will be dependent on the presence of a pool of nuclear RNA to bind the RBPs and increase their dwell time in the nucleus (Fig. 2D).

To test whether nuclear RNA is required for PABP accumulation in the nucleus, we assayed PABP localization when nuclear RNAs are degraded concurrently with cytoplasmic RNAs in response to RNase L activation (Fig 2D). To do this, we used A549 cells that constitutively express RNase L tagged with a nuclear localization signal (NLS) that we previously generated and termed NLS-RNase L (Decker et al., 2021). Unlike RNase L, which strictly localizes to the cytoplasm (Fig. 2C), the RL-NLS localizes to both the cytosol and nucleus (Fig. 2E) and degrades nuclear RNAs in response to poly(I:C) lipofection (Decker et al., 2021). We examined PABP localization via immunofluorescence and RNA levels via poly(dT) FISH post-poly(I:C) lipofection.

An important result is that RNase L-mediated degradation of both nuclear and cytoplasmic RNA reduced the accumulation of PABP in the nucleus. Specifically, in cells where RNase L is targeted to the nucleus (A549-NLS-RL), we observed a reduction in poly(A)+ RNA staining in both the nucleus and cytoplasm following poly(I:C) lipofection in comparison to mock-treated A549 cells (Fig. 2F,G). Importantly, PABP remained predominately localized to the cytoplasm in A549-NLS-RL cells in which RNase L degraded both cytoplasmic and nuclear RNAs (Fig. 2,F,G). In contrast, transfection of poly(I:C) in A549 cells only reduced cytoplasmic poly(A)+ RNA, not nuclear poly(A)+ RNA in comparison to mock treated cells (Fig. 2F,G). In these cells, PABP staining increased in the nucleus and decreased in the cytoplasm in comparison to mock-treated cells (Fig. 2F,G).

These data demonstrate that RNase L activation increases the nuclear localization of several RBPs by increasing the relative number of RBP-binding sites in the nucleus relative to the cytoplasm as a result of degradation of cytoplasmic but not nuclear mRNAs.

### RNase L alters host mRNA processing

The above data imply that RBPs translocated to the nucleus upon RNase L activation associate with RNA. This influx of RBPs would be predicted to compete for RNA-binding with RBPs involved in mRNA processing and thereby alter nuclear RNA processing. To examine if RNase L activation alters RNA processing, we analyzed high-throughput RNA sequencing (RNA-seq) of WT and RL-KO cells following six hours of either mock treatment or poly(I:C) lipofection (Burke et al., 2019).

To examine if there were changes to alternative splicing, we utilized MAJIQ (see methods) which identifies changes in alternative splicing patterns. Splicing changes are expressed for a given splicing event as the Percentage Spliced In (PSI). Thus, we compared the PSI of splicing events between WT or RL KO cells treated with or without poly(I:C) to identify changes in RNA splicing that occur in response to dsRNA and that are dependent on RNase L.

This analysis identified 140 splicing events, across 136 genes, that showed differential splicing either due to poly(I:C) treatment in the WT cells, or were different between the WT and RL KO cell lines post-poly(I:C). These changes are calculated as the difference in PSI between two conditions, the ΔPSI. The changes in splicing in both cases were generally correlated, which indicates that the majority of changes in alternative splicing observed are due to activation of RNase L (Figure 3A). Splicing alteration in 60 genes were statistically significant under both comparisons (Figure 3B). Strikingly, 14 of these 60 genes encode RBPs involved in pre-mRNA splicing. This is notable since many RBPs autoregulate their own splicing (Muller-McNicoll et al., 2019), and this would be consistent with increased nuclear occupancy of RBPs following activation of RNase L.

**Fig. 3.**
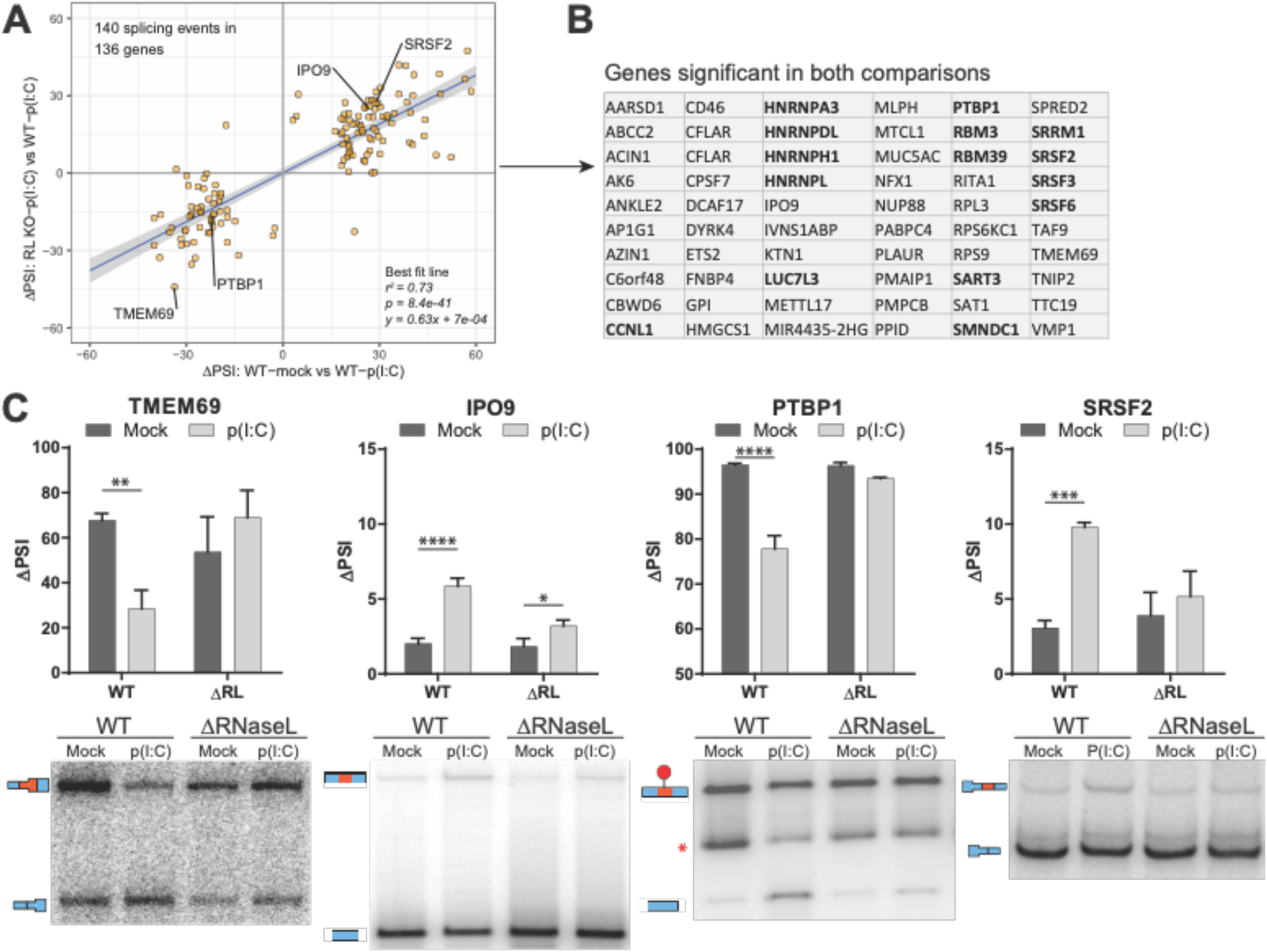
RNase L activation results in alterations to alternative splicing. (A) Scatterplot comparing ΔPSI values in RL KO vs WT cells treated with poly(I:C) on the y axis and WT control vs WT poly(I:C) treated cells on the x axis. The 140 splicing events shown are significant in at least one comparison. Genes validated in panel C are labeled. (B) List of genes significant in both comparisons from panel A. RBPs known to regulate splicing are shown in bold. (C) Validation of splicing events by low cycle radiolabeled RT-PCR. The red asterisk in the PTBP1 gel is a nonspecific product. Bar graphs show quantification of 4 biological replicates. * is p < 0.05, ** is p < 0.01, *** is p < 0.001, *** is p < 0.0001

To validate the analysis of RNA-Seq data, we prepared RNA from mock and poly(I:C) treated WT and RL-KO cells and examined a subset of splicing events by low cycle radiolabeled RT-PCR. In all four cases examined (TMEM69, IPO9, PTBP1, and SRSF2), we observed a changed in exon inclusion with poly(I:C) treatment that was dependent on RNase L (Figure 3C). This demonstrates that RNase L activation can lead to changes in alternative splicing patterns.

Another alteration of splicing can be overall decreased splicing rates and the increased retention of introns (intron retention (IR)). To determine if intron retention is affected by RNase L activation, we calculated an intron/exon ratio (TPM normalized intron over TPM normalized exon counts) per RNA for WT and RL-KO cells with or without poly(I:C) transfection. An increase in intron retention correlates with an increase in the intron/exon ratio.

A striking result was that density plots of all RNAs showed a shift to higher intron/exon ratios upon poly(I:C) treatment in WT cells, highlighting an RNase L-dependent increase in IR (Fig. 4A). The same shift was also observed for transcripts that are either unchanged (Fig. S1A), downregulated (Fig. S1C) or upregulated (Fig. S1E) in WT cells upon poly(I:C) treatment. Consistent with these analyses, IGV traces of multiple example genes displayed increased reads mapping to introns (Fig. 4B, S2B,D,F).

**Fig. 4.**
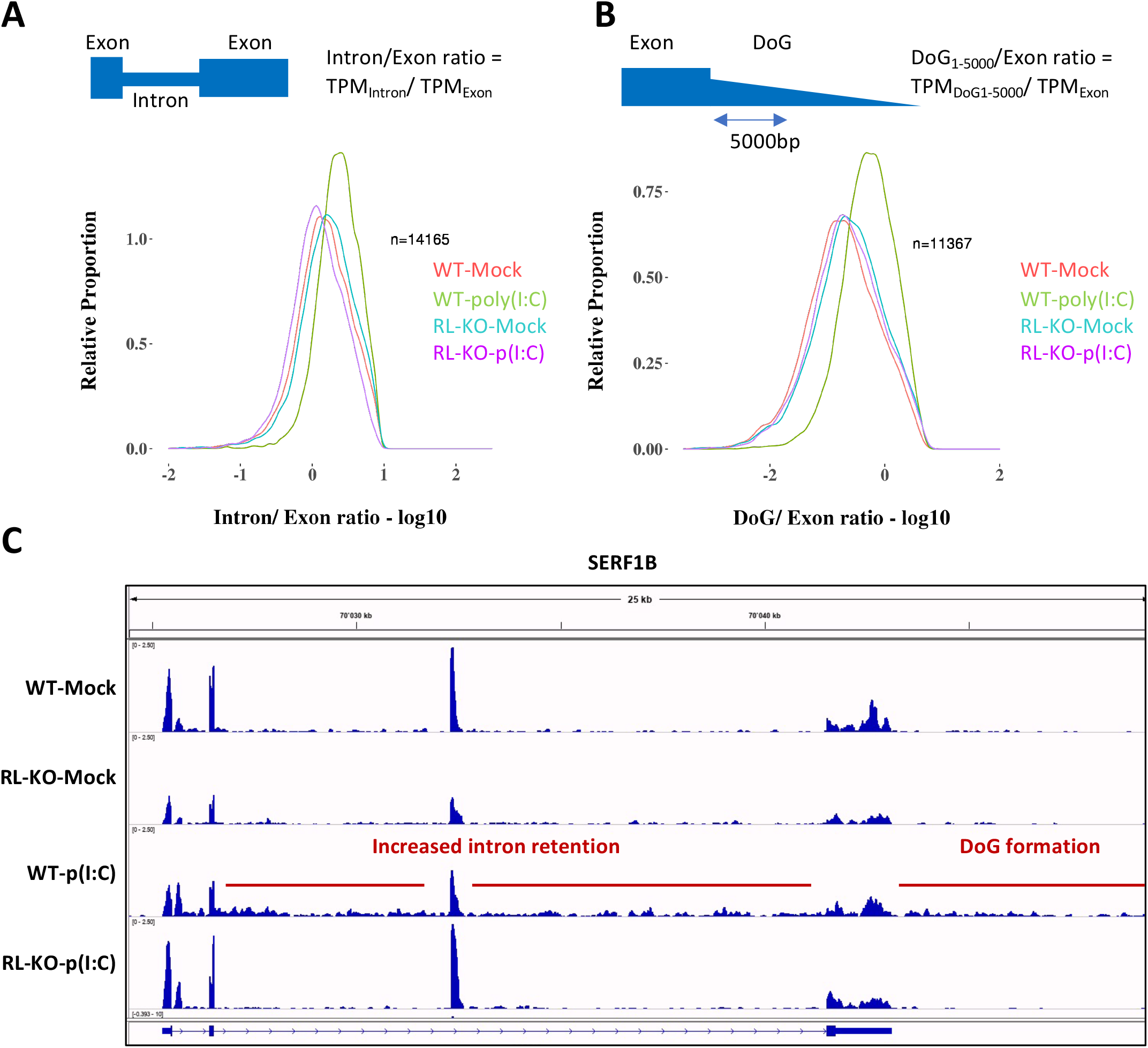
RNase L activation results in global alterations to host RNA processing. (A) Distribution of intron/exon ratios of host RNAs in WT and RL-KO cells following mock or poly(I:C) lipofection. (B) Distribution of DoG_1-5000bp_/exon ratios of host RNAs in WT and RL-KO cells following mock or poly(I:C) lipofection. (C) IGV traces mapping to an example gene. Intron retention and DoG formation is highlighted in WT cells following poly(I:C) lipofection.

We note that for upregulated genes, a shift towards higher intron/exon ratio is visible in wild-type cells compared to RL-KO cells, consistent with an increase in intron retention. However, an additional shift is also visible in both unstressed conditions (Fig. S2C). This is caused by multiple factors. First, a few genes showed high intron retention in RL-KO cells even without stress. Second, transcriptional read-through from upstream genes (see below), causes increased ratios. The majority of such transcripts is filtered out by initial filtering steps (material and methods), however, especially for upregulated genes, some transcripts remain and can cause false positive increased ratios. Third, wrong gene annotations (e.g. an exon with increased reads is counted within intronic region), can make increase in ratio for upregulated genes more pronounced. And lastly, despite an overall increase in IR in WT cells upon poly(I:C) (Fig. S1F), the intron/exon ratio decreases due to the increased expression and higher exon counts over the intron counts, especially for genes with short introns and large exons.

These data argue that RNase L activation also decreases the efficiency of intron removal, which is supported by smFISH data of selected targets (see below).

We also observed that RNase L activation perturbs transcription termination. Specifically, we examined whether RNase L activation resulted in downstream of gene (DoG) transcriptional read-through, which are observed during viral infection and various other stresses and are consistent with defects in transcription termination (Vilborg and Steitz, 2017; Vilborg et al., 2015; Rutkowski et al., 2015; Bauer et al., 2018). To do this, we calculated an DoG1-5000/exon ratio, where DoG1-5000 corresponds to TPM normalized counts over the first 5000 nt after the annotated gene end (Fig. 4B and Fig. S2A,C,D) (see methods).

In WT cells transfected with poly(I:C), we observed DoG transcripts, which were notably absent in poly(I:C)-transfected RL-KO cells (Fig. 4B,C). For example, IGV traces for multiple RNAs demonstrates increased reads mapping to the DoG region in WT cells treated with poly(I:C) that are absent in mock-treated WT or RL-KO cells as well as poly(I:C)-treated RL-KO cells (Fig. 4C and Fig. S2,B,D,F).

We confirmed DoG formation in the long non-coding RNA, NORAD, using two smFISH probes, with one targeting the NORAD RNA region and a second targeting the DoG region of NORAD (Fig. S3A). Staining cells twelve hours post-poly(I:C) lipofection revealed accumulation of RNAs with 3’ DoGs as detected by hybridization to the probe downstream of the NORAD coding region in WT but not RL-KO cells (Fig. S3B). Consistent with earlier results, we also observed RNase L-mediated degradation of the cytosolic NORAD RNA (Burke et al., 2019), and the accumulation of NORAD RNA in the nucleus due to a block to nuclear export (Burke et al., 2021). As expected, in the RL-KO cells, cytosolic NORAD RNA was not degraded and accumulated in stress granules (Khong et al., 2017). DoG RNAs co-localized with NORAD RNA at both the site of transcription (2 large/intense foci) as well as dispersed in the nucleus as individual RNAs (Fig. S3B).

These observations indicate that RNase L activation results in global nuclear RNA processing alterations, including changes in alternative splicing, intron retention and reduced transcription termination.

### RNase L activation alters antiviral mRNA biogenesis

A key aspect of the cellular response to dsRNA is the induction of antiviral gene expression. Given this, we examined how RNase L activity affected the processing of mRNAs transcriptionally induced by dsRNA, including type I and type III interferons (IFNs). We observed that RNase L altered the RNA processing of the pre-mRNAs for both types of interferons. For example, we observed increased RNA-seq reads mapping downstream of *IFNB1* and *IFNL1* in WT cells in comparison to RL-KO cells (Fig. 5A,B), indicating downstream of gene (DoG) transcriptional read-through when RNase L is activated. Moreover, we observed increased reads mapping to the introns of IFNL1, indicative of intron retention (Fig. 5B). To validate these data, we performed smFISH using probes that target the DoG of IFNB1 and IFNL1 or the introns of IFNL1.

**Fig. 5.**
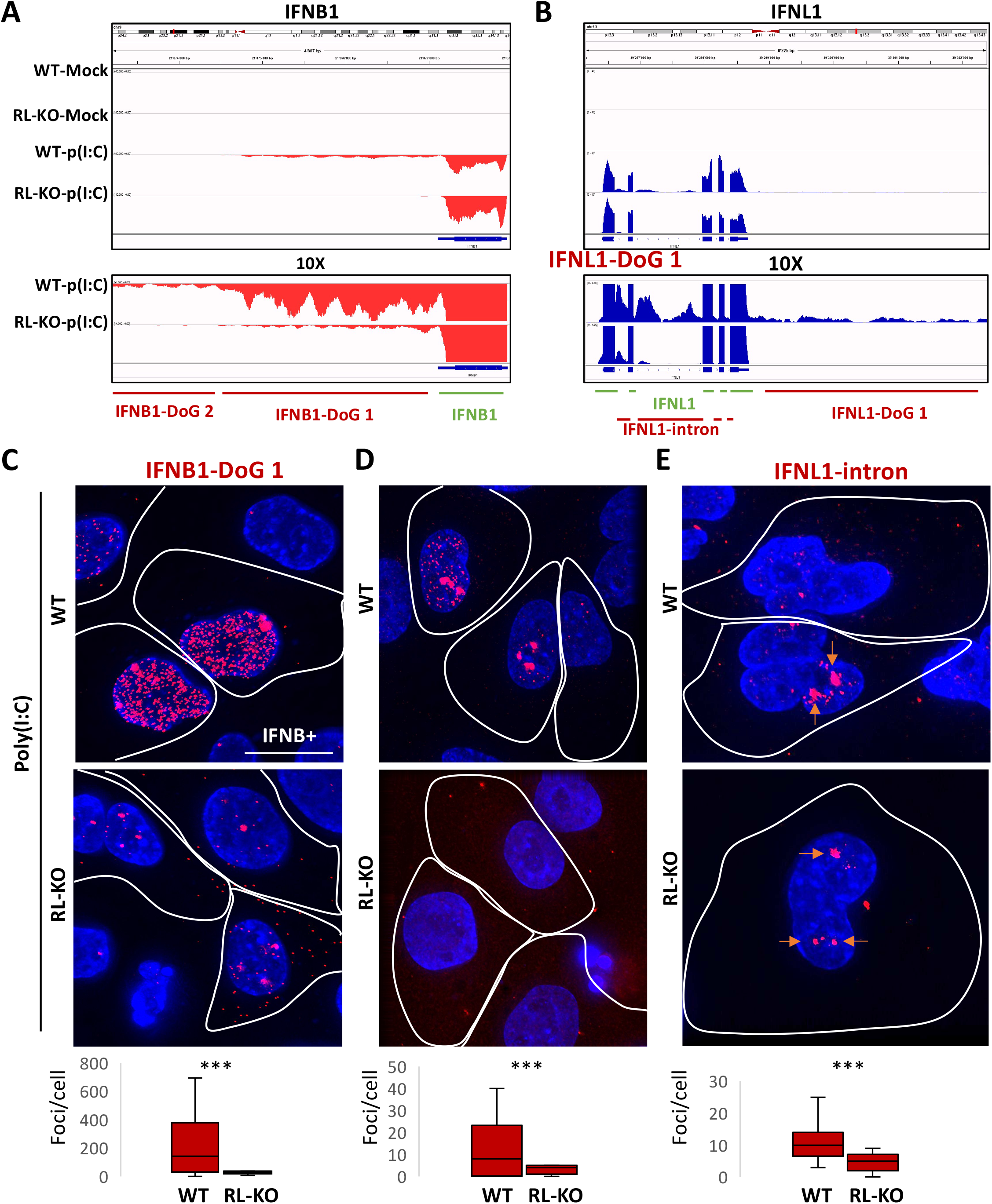
RNase L promotes DoG transcriptional read-through and intron retention in type I and type III interferon RNAs. (A) IGV traces mapping to IFNB1. Below shows the regions targeted by smFISH probes. (B) IGV traces mapping to IFNL1. Below shows the regions targeted by smFISH probes. (C) smFISH for IFNB1-DoG sixteen hours post-lipofection of poly(I:C) in WT and RL-KO cells. The cells that induced IFNB1, as determined by smFISH for the CDS of IFNB1 (Fig. S4A,B), are demarcated by a white line. IFNB1 DoG smFISH foci are quantified in WT an RL-KO cells in the graph below. (D and E) similar to (C) but for (D) IFNL1-DoG-1 RNA sixteen hours post-lipofection of poly(I:C) or (E) IFNL1-intron RNA twelve hours post-lipofection of poly(I:C). Staining and quantification of IFNL1 CDS is shown in Fig. S4C,D,E.

Consistent with our RNA-seq analyses, several observations indicate that RNase L promotes transcriptional read-through of *IFNL1* and *IFNB1* genes. First, in a portion of WT cells that induced IFNB1 or IFNL1 expression in response to poly(I:C), as determined by co-smFISH for the CDS regions of these genes (Fig. S4A,B,C,D,E), we observed abundant and disseminated DoG smFISH foci (Fig. 5B,C,D). We note that we did not observe abundant smFISH foci targeting the DoG-2 region, which is further downstream of the DoG 1 region of IFNB in WT cells (Fig S5A). This correlates with lower reads mapping to this region and consistent with lower abundance of these transcripts as assessed by RNA-seq. We did observe staining for the DoG-2 region of IFNL1, but note that it was predominantly localized to the sites of transcription (Fig. S5B).

*IFNB1* and *IFNL1* DoG RNA staining was significantly less abundant in RL-KO cells in comparison to WT cells despite RL-KO cells containing higher levels of CDS staining for both IFNB1 and IFNL1 (Fig. S4A,B,C,D). Notably, IFNB1-DoG staining was mostly confined to large foci that are consistent with IFNB1 sites of transcription in RL-KO cell (Fig. 5C), and the IFNL1-DoG RNA was not observed in RL-KO cells (Fig. 5D). These data demonstrate that that RNase L promotes DoG read-through transcription of *IFNB1* and *IFNL1* genes.

*IFNL1* intron staining was largely observed at sites consistent with the IFNL1 sites of transcription, and this was observed in both WT and RNase L-KO cells (Fig. 5E). Nevertheless, we observed increased IFNL1-intron smFISH foci in WT cells in comparison to RL-KO cells (Fig. 5E). This observation is consistent with the RNA-seq data and demonstrates that RNase L activation promotes intron retention in *IFNL* transcripts.

### Alterations in RNA processing contribute to nuclear retention of antiviral mRNAs

We have previously documented that RNase L activation can lead to nuclear retention of host and viral mRNAs (Burke et al., 2021a). The observation that IFNB1 and IFNL1 DoG RNAs are largely localized to the nucleus led us to examine whether DoG RNA included on the IFNB1 or IFNL1 transcripts could contribute to their nuclear retention. To examine this, we analyzed the localization of DoG RNA and CDS RNA in WT and RL-KO cells following poly(I:C) lipofection.

Several observations indicate that DoG production from the IFNB1 mRNA contributes to its nuclear retention. First, the IFNB1-DoG RNAs are almost exclusively localized in the nucleus, even in cells that have exported IFNB mRNAs that only stain for the CDS region to the cytosol (Fig. 6A,B; red arrows). Second, in WT cells with abundant and disseminated IFNB1-DoG foci, the DoG foci co-localize with IFNB1-CDS foci that are retained in the nucleus (Fig. 6A; red arrows). Third, IFNB1-CDS foci located in the cytoplasm only contain DoGs in very rare cases (Fig. 6A; red arrows). Lastly, we observed that the abundance of DoG foci positively correlates with the ratio (nuclear/total) of IFNB-CDS or the absolute number of nuclear IFNB-CDS foci (Fig. 6C,D). We observed similar effects eight hours post-transfection with poly (I:C) (Fig. S6A,B,C,D,E). These data argue that IFNB1 RNA transcripts containing the DoG RNA are not exported to the cytoplasm and argue that DoG formation contributes to the inhibition of export of IFNB mRNAs.

**Fig. 6.**
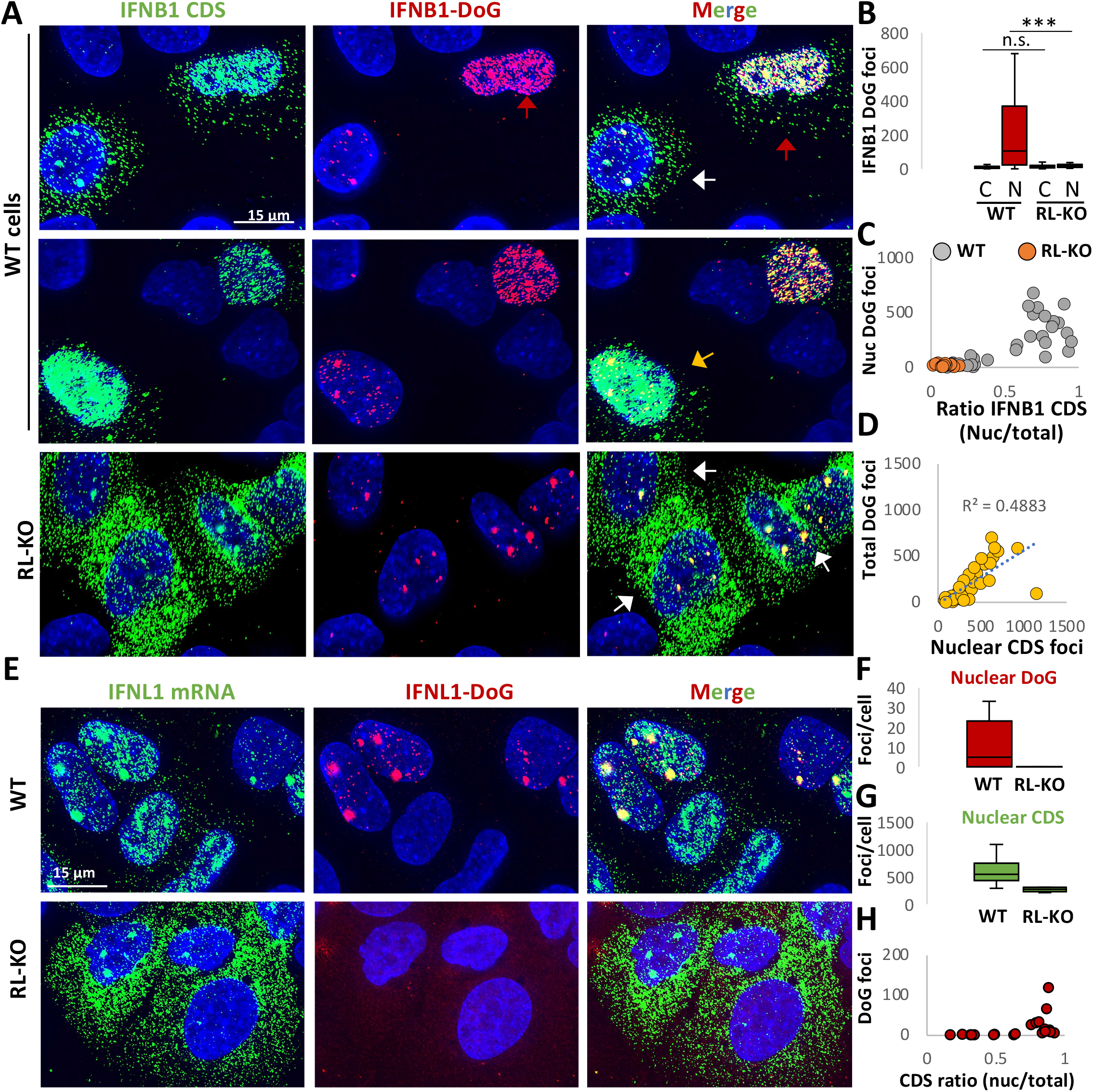
DoG RNA included on interferon-encoding mRNAs correlates with their nuclear retention. (A) Co-smFISH for the CDS and DoG-1 regions of IFNB1 sixteen hours post-lipofection of poly(I:C). (B) Box plots displaying of the number of IFNB1-DoG foci localized to the nucleus (n) or cytoplasm (c) in WT or RL-KO cells as represented in (A). (C) Scatter plot of the ratio (nucleus/cytoplasm) of IFNB1-CDS foci (x-axis) and nuclear IFNB1-DoG foci (y-axis) show positive correlation between DoG RNA and nuclear retention. (D) Scatter plots of the quantity of nuclear IFNB1-DoG foci (y-axis) and the quantity of nuclear IFNB1-CDS foci shows IFNB1 DoG RNA increase as the absolute number of IFNB1 CDS smFISH in the nucleus increases. (E) Co-smFISH for the CDS and DoG-1 regions of IFNL1 sixteen hours post-lipofection of poly(I:C). (F and G) Quantification of (F) nuclear IFNL1-DoG-1 foci or (G) IFNL1-CDS foci as represented in (E). (H) Scatterplot of the ratio (nucleus/cytoplasm) of IFNL1-CDS foci (x-axis) and nuclear IFNL1-DoG foci (y-axis).

Our data suggests that additional mechanisms can inhibit mRNA export. For IFNB mRNA, we also observed cells in which most nuclear-retained IFNB1-CDS RNAs did not contain IFNB-DoG RNA (Fig. 6A, yellow arrow). The accumulation of IFNB mRNAs in the nucleus that do not hybridize to DoG probes suggests two possibilities. First, there could be a mechanism independent of DoG transcriptional read-through that inhibits *IFNB1* mRNA export, which is supported by our analysis of IFNL1 mRNA (see below). However, we cannot rule out the formal possibility that all the nuclear retained IFNB RNAs could contain DoGs, but with some being too short to hybridize to the DoG probes. However, since the vast majority of RNA-seq reads end at the normal 3’ end of IFNB mRNAs (Fig. 5A), we consider this possibility unlikely.

The examination of IFNL1 mRNAs provides additional evidence for a block to mRNA export that is independent of RNA processing defects. Specifically, we observed that most of the nuclear-retained IFNL1 mRNA did not hybridize to smFISH probes for IFN1L introns or DoGs and were much more abundant than the DoG and intron foci (Fig. 6E,F,G). We note that we did observe that IFN1L mRNAs that contained DoGs or intron sequences were mostly nuclear-retained (Fig. 6E,H), consistent with defects in RNA processing limiting RNA export.

Taken together, these observations demonstrate the defects in RNA processing can contribute to nuclear retention of mRNAs after RNase L activation, but also provide evidence for a block to mRNA export independent of DoG-RNA inclusion and intron retention.

### Re-localization of RBPs and RNA processing alterations occur during viral infection

One limitation of the above experiments is that we have used poly(I:C) as a dsRNA mimic of viral infection. In order to verify that these alterations were relevant to viral infection, we examined whether RNase L-dependent re-localization of RBPs and nuclear RNA processing alterations occurred during viral infection. We analyzed PABP localization via immunofluorescence in WT and RL-KO A549 cells following infection with either dengue virus serotype 2 (DENV2) or SARS-CoV-2 (using A549^ACE2^ cells), both of which activate RNase L (Burke et al., 2021a; Burke et al., 2021b; Li et al., 2021). To identify infected cells, we co-stained the viral mRNAs via single-molecule fluorescence in situ hybridization (smFISH), while alterations in RNA processing were examined by smFISH for IFNB of IFN1L DoGs or introns.

These analyses showed that RNase L activation promotes PABP translocation to the nucleus and alteration of RNA processing in response to SARS-CoV-2 infection (Fig. 7A,B,C,D,E). Specifically, all WT A549 cells infected with SARS-CoV-2, as detected by smFISH for the viral RNA, displayed accumulation of PABP in the nucleus (Figure 7A; white arrows), which was not observed in RL-KO cells. Moreover, smFISH for IFNB1 CDS and DoG demonstrated the production of IFNB1 DoGs during SARS-CoV-2 infection was higher in WT but not RL-KO cells (Fig. 7 D,E). Notably, IFNB1 transcripts containing DoG RNAs were largely localized in the nucleus (Fig. 7D).

**Fig. 7.**
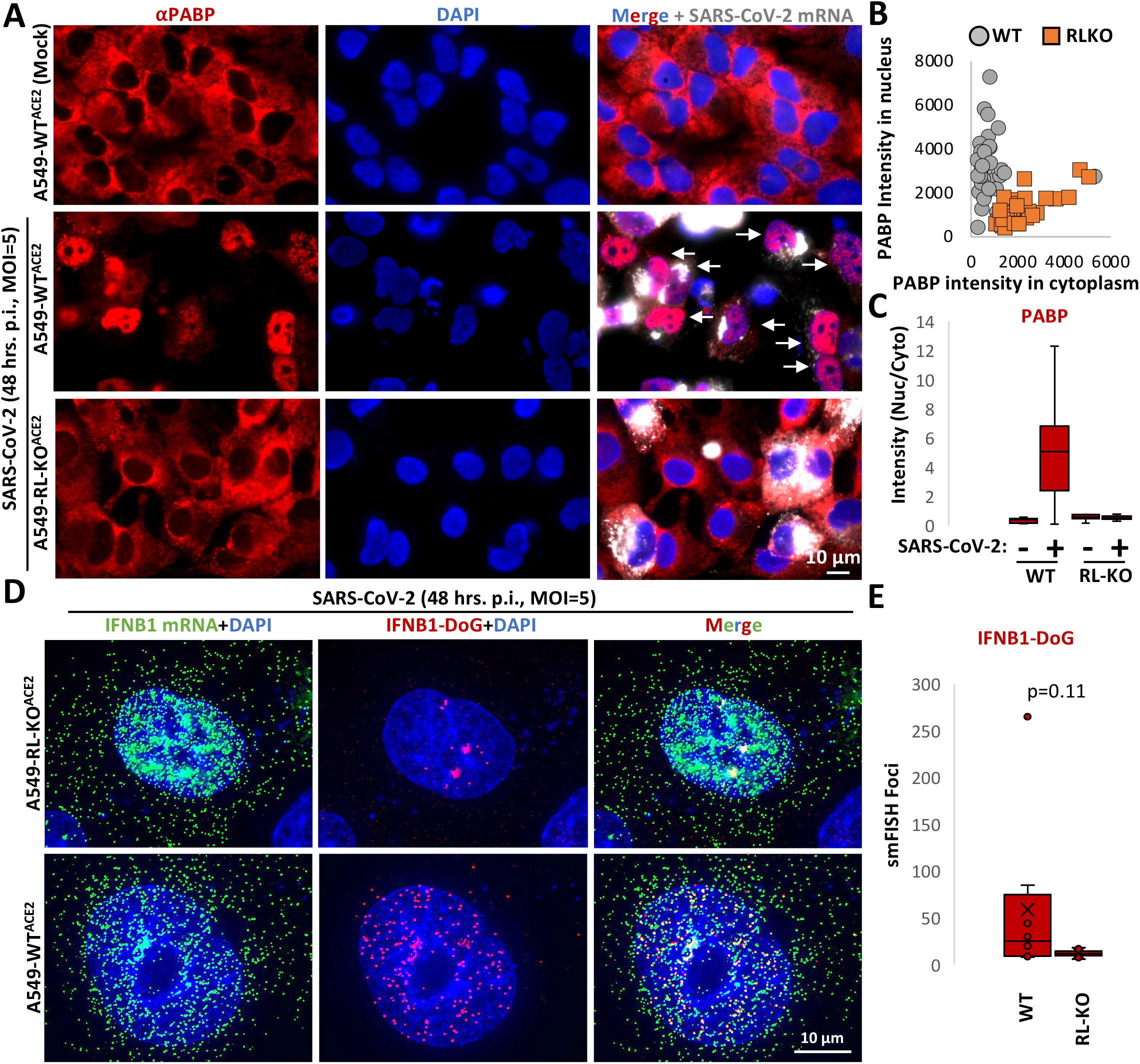
RNase L promotes nuclear RBP influx and DoG transcriptional read-through of IFNB1 during SARS-CoV-2 infection. (A) Immunofluorescence assay for PABP in WT^ACE2^ and RL-KO^ACE2^ A549 cells forty-eight hours post-infection with SARS-CoV-2 (MOI=5) or mock-infected WT cells. To identify infected cells, smFISH for SARS-CoV-2 ORF1b mRNA was performed. (B) Scatter plot of mean intensity values for PABP staining in the nucleus (y-axis) or cytoplasm (x-axis) in mock or SARS-CoV-2-infected WT^ACE2^ or RL-KO^ACE2^ A549 cells as represented in (A). Dots represent individual cells. (C) Box plot of the ratio (nucleus/cytoplasm) of the mean intensity of PABP as represented in (A). (D) smFISH for IFNB1-CDS and IFNB1-DoG forty-eight hours post-infection with SARS-CoV-2 in WT^ACE2^ or RL-KO^ACE2^ A549 cells (MOI=5). (E) Quantification of IFNB1 DoG RNA in WT and RL-KO cells.

Similar results were observed with DENV-infected cells, although approximately half of DENV-infected cells do not activate RNase L-mediated mRNA decay (Burke et al., 2021a) (Fig. S7A). Thus, approximately half of DENV-infected cells did not display translocation of PABP to the nucleus as expected (Fig. 8A and Fig. S7B). However, we observed many DENV2-infected WT cells assembled RLBs, whereas many DENV2-infected RL-KO cells assembled SGs (Fig. 8A). This allowed us to identify cells that activated the dsRNA immune response. We then calculated the nuclear to cytoplasmic PABP in DENV-infected WT cells that activated RNase L (RLB+) or RL-KO cells that activated PKR (SG+) in comparison to cells that did not activate dsRNA response (RL- for WT cells or SG- for RL-KO cells). These analyses revealed a substantial increase in nuclear PABP specifically in DENV2-infected WT cells that activated RNase L (Fig. 8B,C).

**Fig. 8.**
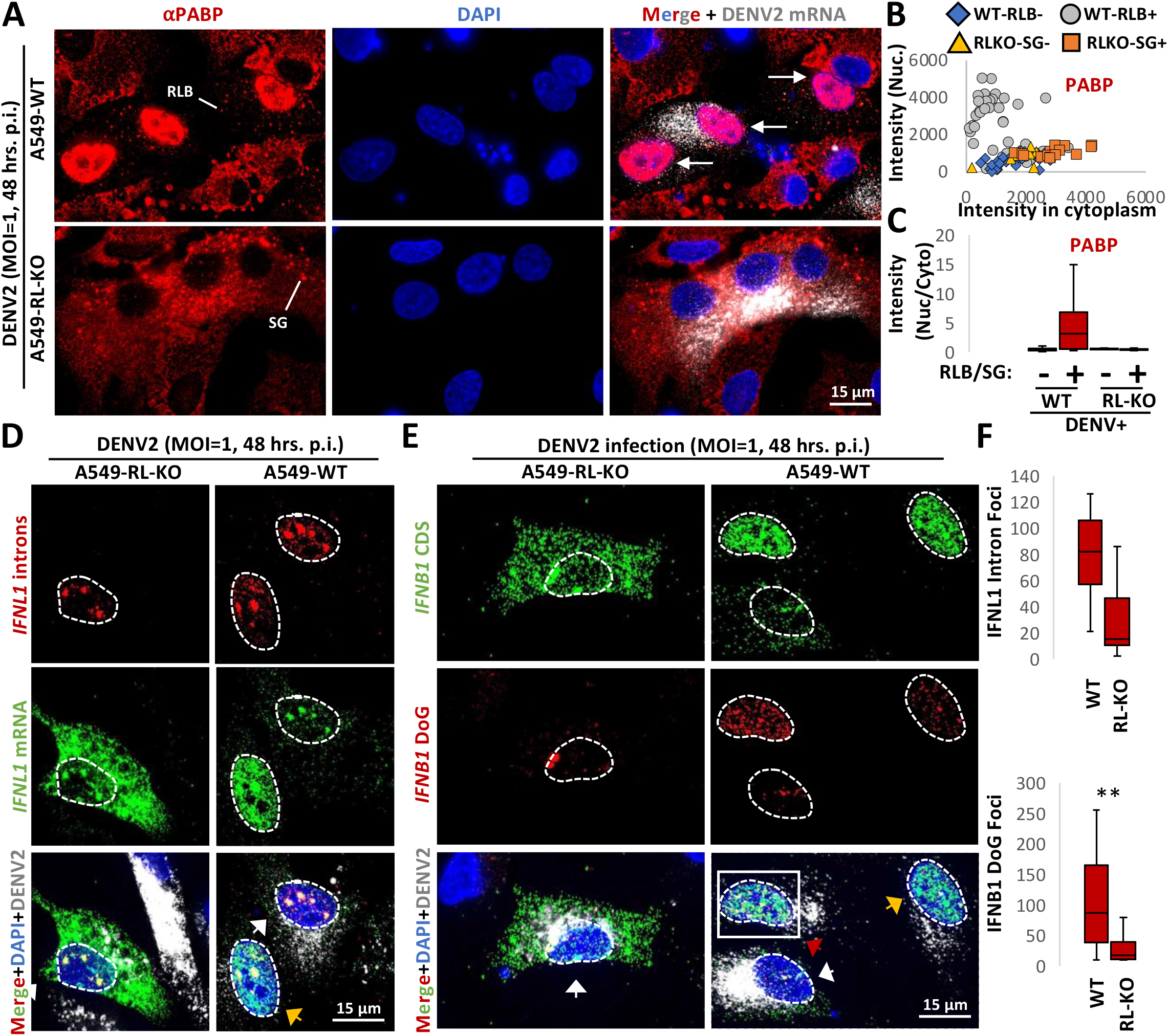
RNase L promotes nuclear RBP influx and DoG transcriptional read-through and intron retention during dengue virus infection. (A) Immunofluorescence assay for PABP in WT and RL-KO A549 cells infected with dengue virus serotype 2 (DENV2) forty-eight hours post-infection (MOI=1). smFISH for DENV2 mRNA was performed to identify infected cells. (B) Scatter plot of mean intensity values for PABP staining in WT or RL-KO cells that did or did not activate the dsRNA response based on RLB assembly (WT cells) or SG assembly (RL-KO) cells as represented in (A). (C) Box plot of the ratio (nucleus/cytoplasm) of the mean intensity of PABP (B). (D) smFISH for IFNL1-CDS, IFNL1-intron, and DENV mRNA in WT or RL-KO cells forty-eight hours post-infection with DENV2 (MOI=1). (E) smFISH for IFNB1-CDS and IFNB1-DoG in WT or RL-KO cells forty-eight hours post-infection with DENV2 (MOI=1). (F) Box plots quantifying the smFISH from (D and E).

Examination of RNA processing defects in DENV-infected cells via smFISH revealed that RNase L-dependent RNA processing alterations occurred during these infections. Specifically, we observed that IFNL1-intron and IFNB1-DoG were both higher in DENV-infected WT cells comparison to RL-KO cells (Fig. 8D,E,F).

Taken together, the analysis of IFN mRNAs documents that RNase L activation either due to poly(I:C) transfection or viral infection triggers the accumulation of RBPs in the nucleus and affects nuclear RNA processing of antiviral mRNAs, with a stronger effect on transcriptional termination leading to the production of DoGs for these mRNAs.

## DISCUSSION

Several observations support that RNase L-mediated RNA decay results in re-localization of RBPs from the cytoplasm to the nucleus, which in turn alters nuclear RNA processing (Fig. 9). First, we observed that several RBPs concentrate in the nucleus following RNase L activation (Fig. 1A,B). Second, RNase L and RNase L-mediated RNA decay are localized the cytoplasm (Fig. 2A,B,C), sparing nuclear RNA from degradation. Moreover, intact nuclear RNA is required for the accumulation of RBPs to the nucleus, suggesting RBPs associate with nuclear RNA upon influx into the nucleus (Fig. 2E,F,G). Third, our RNA-seq analyses show that RNase L activation triggers global alterations in splicing and transcription termination (Figs. 3 and 4). Fourth, our smFISH analyses show that the number of DoGs and introns are increased in WT cells in comparison to RL-KO cells in response to poly(I:C) lipofection, and that these RNAs can be retained at the sites of transcription or disseminated in the nucleus (Fig. 5C,D,E, Fig. 6A,E, Fig. S3,B). Importantly, we show that RNase L-dependent RBP re-localization to the nucleus and alterations to type I and type III interferon mRNA biogenesis occur during dengue virus or SARS-CoV-2 infection (Figs. 7 and 8).

**Fig. 9.**
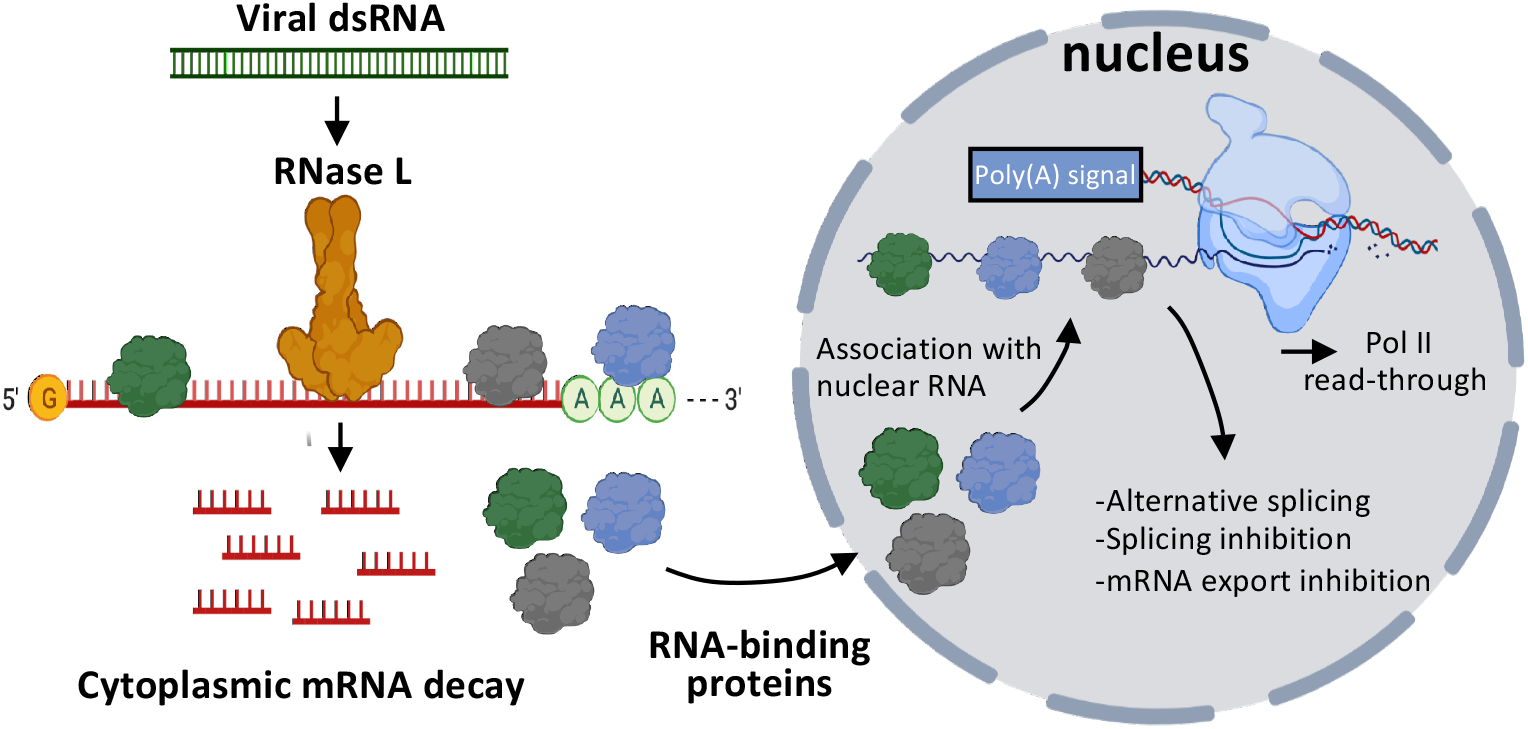
Model for RNase L-mediated regulation of nuclear RNA processing. Activation of RNase L in response to viral double-stranded RNA (dsRNA) results decay of cytoplasmic RNAs. RNA-binding proteins (RBPs) disassociate from degraded cytoplasmic RNA and shuttle to the nucleus where they bind to nuclear RNA. The binding of RBPs to nuclear RNA alters RNA processing events.

This work highlights the fundamental conclusion that the availability of RNA-binding sites can influence the distribution of RBPs between the cytosol and nucleus, which explains multiple observations in the literature. First, during the integrated stress response, the release of mRNAs from ribosomes due to phosphorylation of eIF2α exposes new RNA-binding sites for RBPs in the cytoplasm, which causes nuclear RBPs to re-distribute to the cytoplasm and/or stress granules (Fig. 1C, CUGBP1 and TIA-1) (Kedersha et al., 1999). Similarly, the export of RBPs from the nucleus in response to actinomycin D-mediated inhibition of transcription can be understood as a loss of nuclear RNA leading RBPs to associate with cytosolic RNA (Cáceres et al., 1998). Finally, when cytosolic RNAs are degraded by the KSHV viral Sox2 nuclease, numerous RBPs are observed to accumulate in the nucleus (Gilbertson et al., 2018). Thus, sub-cellular changes in RNA abundance and/or availability of RNA binding sites dictates RBP localization for many RBPs. Indeed, a number of RBPs known to act in mRNA processing are themselves differentially spliced upon exposure to poly(I:C) in an RNase L-dependent manner (Fig 3). Interestingly, some RBPs are less affected by cytosolic RNA decay with respect to their localization. For example, G3BP1 shows only a small increase in nuclear accumulation following RNase L activation (Fig. 1B,C). We suspect that this is due to their association with protein substrates that regulate their localization, or a slower intrinsic rate of protein import into the nucleus.

RNase L-mediated regulation of RBP localization represents a new and important potential antiviral mechanism since studies have shown that host RBPs interact with and can affect viral replication (Li and Nagy, 2011). Since viruses replicate specifically in either the nucleus or cytoplasm, the RNase L-mediated re-localization of RBPs between the cytoplasm and nucleus may promote the antiviral function of RBPs and/or viruses may use RNA decay to avoid host RBPs. Notably, several viruses, including Kaposi’s sarcoma-associated herpesvirus (KSHV) and herpes simplex virus 1 (HSV-1) encode for proteins (SOX, VHS, respectively) that degrade cellular RNAs, inhibit nuclear mRNA export, and cause nuclear RNA-binding protein influx (Glaunsinger et al., 2005; Kumar and Glaunsinger, 2010; Elliot et al., 2018). Interestingly, KSHV encodes an RNA, PAN, that rescues nuclear export of viral mRNAs (Withers et al., 2018). An intriguing possibility is that the PAN RNA titrates RBPs that interfere with nuclear export, thereby allowing export of the herpes mRNAs. Thus, RBP re-localization can be either host- or viral-mediated and can potentially alter both host and viral gene expression. Lastly, the inhibition of host-mediated RNA decay by viruses, such as by flavivirus sfRNA-mediated inhibition of XRN-1 or poliovirus-mediated inhibition of RNase L (Moon et al., 2012; Han et al., 2007), may regulate host RBP interactions with the viral RNA. Elucidating how overall RNA abundance and subcellular location regulates the availability RBPs to modulate both host and viral RNA function will be an important issue to address.

An interesting observation is that SARS-CoV-2 infection, which results in rapid degradation of host mRNAs even in the absence of RNase L through the action of the SAR2 Nsp1 protein (Burke et al., 2021b), does not trigger robust PABP translocation to the nucleus as demonstrated by the lack of nuclear PABP staining in SARS-CoV-2 infected RL-KO cells (Fig 7B,D,E). We suggest two possibilities to explain this phenomenon. First, the difference in the RNA decay mechanisms between RNase L and SARS-CoV-2 Nsp1 could result in differential release of RBPs such as PABP from RNAs. Second, SARS-CoV-2 may prevent RBP influx into the nucleus, and RNase L overcomes this inhibitory mechanism. Nevertheless, during SARS-CoV-2 infection we still observed more IFNB1 DoG RNAs in WT cells than RL-KO cells, indicating that DoG generation is a consequence of RNase L activation in response to SARS-CoV-2 infection. Whether this is beneficial for SARS-CoV-2 since it reduces host antiviral gene expression is unclear. However, since RNase L only reduces SARS-CoV-2 replication via decay of the viral mRNA by only 3-4 fold (Li et al., 2021; Burke et al., 2021), perhaps the potential loss of fitness of allowing RNase L activation is outweighed by the perturbation in host interferon production.

Our data establish that RNase L activation promotes the formation of DoG transcription read-through. Interestingly, while DoG transcripts are generally retained at the site of transcription (Vilborg et al, 2015), we observed abundant IFNB1 DoG transcripts that contained the IFNB1 CDS disseminated throughout the nucleus (Fig. 6A,B). Moreover, the IFNB1 transcripts with DoG RNA were almost exclusively localized to the nucleus, whereas IFNB1 transcripts in the same cells without the DoG RNA were localized to the cytoplasm. These observations suggest that the DoG RNA on IFNB1 transcripts inhibits their mRNA export, even after their release from the site of transcription. The inclusion of downstream elements on the IFNB1 mRNA transcript may present a new RNase L-dependent regulatory mechanism, which will be a focus of future work.

While much lower in abundance and more localized to the IFNB1 transcription site in comparison to WT cells, the IFNB1 DoG also formed in RL-KO cells (Fig. 5B). This indicates that IFNB1 DoG formation is a normal aspect of IFNB1 gene induction. However, unlike IFNB1, we did not observe any IFNL1 DoG RNA in RL-KO cells. Thus, DoG transcriptional read-through is differential with respect to different dsRNA-induced genes. Understanding this difference may reveal key aspects for transcriptional and RNA processing regulatory mechanisms of these genes.

The RNA processing defects promoted by RNase L activation, such as DoG read-through transcription and splicing alterations (alternative splicing and intron retention), may prove to be new antiviral mechanisms that perturb viral gene expression or promote the host immune response. For example, DoG transcriptional read-through of dsDNA viral genomes could interfere with productive viral gene expression and/or generate PAMPs that could trigger antiviral PRRs. Moreover, since RBPs can directly modulate transcription (Xiao et al., 2019), this work strongly implies that widespread RNA degradation in the cytosol will lead to changes in transcription of multiple genes due to the influx of RBPs into the nucleus. Future work will examine these potential functions and address the specific mechanism by which RNase L activation alters nuclear RNA processing and transcription.

## ACKNOWLEDGMENTS

Research reported in this publication was supported by the National Institute of Allergy and Infectious Diseases of the National Institutes of Health under Award Number F32AI145112 (J.M.B), funds from HHMI (Roy Parker), National Institutes of Health R01-AI-137011 (S.L.S.), and support provided by the Office of the Vice President for Research and the Dept. of Microbiology, Immunology and Pathology at Colorado State University (Rushika Perera),

## AUTHOR CONTRIBUTIONS

J.M.B. and Roy Parker conceived the project. J.M.B. performed poly(I:C) transfections, western blots, microscopy, and quantification of microscopy. N.R. performed RNA-seq analyses. M.B.F performed splicing analyses and validation. E.R.W.S. performed DENV infections. L.A.S. performed SARS-CoV-2 infections. J.M.B., N.R., and Roy Parker interpreted the data. J.M.B., N.R., and Roy Parker wrote the manuscript.

## MATERIALS & METHODS

### Cell culture

Parental and RNase L knockout A549 cell lines are described in Burke et al., 2019. Cells were maintained at 5% CO_2_ and 37 degrees Celsius in Dulbecco’s modified eagle’ medium (DMEM) supplemented with fetal bovine serum (FBS; 10% v/v) and penicillin/streptomycin (1% v/v). Routine testing for mycoplasma contamination was performed by the cell culture core facility. African green monkey kidney cells (Vero E6, ATCC CRL-1586) were maintained at 5% CO_2_ and 37 degrees Celsius in DMEM supplemented with FBS (10% v/v), 2 mM non-essential amino acids, 2 mM l-glutamine, and 25 mM HEPES buffer.

For transfections, high-molecular weight poly(I:C) (InvivoGen: tlrl-pic) and 3-μl of lipofectamine 2000 (Thermo Fisher Scientific) per 1-ug or poly(I:C) was used per manufacturer’s instructions.

### Viral infections

Viral infections are described in Burke et al., 2021a and Burke et al., 2021b. Briefly, cells were infected with DENV serotype 2 16681 strain at MOI of 1.0. Cells were fixed 48 hours after infection. SARS-CoV-2/WA/20/01 (GenBank MT020880; BEI Resources: NR-52881) was passaged in Vero E6 cells, and viral titer was determined via plaque assay on Vero E6 as previously described in (Dulbecco et al., 1953). A multiplicity of infection (MOI) of 5 was used. All SARS-CoV-2 infections were conducted under biosafety level 3 conditions at Colorado State University. Dengue virus infections were conducted under biosafety level 3 conditions at University of Colorado, Boulder. For infections, cells were seeded in 6-wells format onto cover slips and inoculated twenty-four hours later. Cells were fixed in 4% paraformaldehyde and phosphate-buffered saline (PBS) for 20 minutes, followed by three five-minute washes with 1X PBS, and stored in 75% ethanol.

### RNA-seq analysis

Our previously published RNA-seq data (Burke et al., 2019) were re-processed using a Nextflow pipeline (https://github.com/Dowell-Lab/RNAseq-Flow). Normalized TDF files that were generated by the pipeline were used to visualize representative gene traces with the Integrative Genomics Viewer (Robinson et al., 2012). Exon or intron reads were counted for each sorted BAM file over annotated genes in the RefSeq hg38 genome using featureCounts (Rsubread/2.0.1,(Liao et al., 2013). GTFtools was used to determine intron boundaries (Hong-Dong Li e al., 2018) and to generate an introns only gtf file. RNA isoforms were filtered for the highest expressed isoform equivalent to the highest FPKM value in the WT condition. DESeq2/1.26.0 was used for differential expression analysis. ERCC RNA Spike-Ins were used to estimate and correct size factors in DESeq2. In particular, ERCC Spike-Ins reads were mapped using the same Nextflow pipeline to the ERCC sequences (https://tools.thermofisher.com/content/sfs/manuals/ERCC92.zip), counted using featureCounts and the ERCC Spike-Ins size factors derived by DESeq2. Significant differentially upregulated or downregulated RNAs were selected by baseMean >5, padj < 0.05 and a log2fold change of 1. Nonsignificant genes were selected by padj > 0.05 and log2fold change of −0.5 to 0.5.

Intron retention analysis was performed on all RNAs with the Deseq2 baseMean cutoff > 30, to remove RNAs with too low counts. Intron/exon ratios were derived by diving the TPM normalized intron counts by TPM normalized exon counts. Intron/exon ratios > 8.5 were a result from “towers” in intronic regions, counts in non-annotated regions, noise due to low counts or read-through from an upstream gene. Therefore, all intron/exon ratios >8.5 were removed from analysis.

DoG formation was estimated similarly, by generating TPM normalized counts over the first 5000 pb following the exon annotation. These TPM normalized DoG1-5000 counts were divided by the TPM normalized exon counts to derive and a DoG/exon ratio. For the same reasons as describe for intron/exon ratios, DoG/exon ratios > 5 were removed from analysis. Moreover, small RNAs (exon size < 300bp) were removed from analysis.

In our ratio analysis, genes were not filtered for “clean” genes, genes that have counts due to expression and not due to read-through transcription from up-coming gene (Rosa-Mercado et al., 2021). However, many genes that were not expression but showed read-through transcription from up-coming gene were filtered out by the intron/exon ratios >15 and DoG/exon ratios > 5 step. We do anticipate false-positives left in our analysis, especially in the upregulated group.

Figures were made using ggplot2/3.3.3.

For the splicing analysis, raw fastq reads were trimmed to remove adapters and low quality reads with bbduk/38.79 (Bushnell, 2014), aligned to the hg38 genome using STAR/2.5.2a (Dobin et al., 2013), sorted and indexed with samtools/1.9 (Li et al., 2009), and then analyzed with MAJIQ/2.0 (Vaquero-Garcia et al., 2016). Splicing events were quantified for downstream analysis using the voila classify function, which groups splicing events into distinct modules. Events were considered significant if the probability of having a ΔPSI > 10 was greater than 90%.

### RT-PCR analysis of splicing

Validation of splicing events by low-cycle radiolabeled PCR was performed as previously described (Lynch and Weiss, 2000) with the following modifications. Annealing temperatures during PCR amplification, and the number of cycles required to maintain the signal in the linear range, were determined empirically for each splicing event. Gene specific primers are listed in Table S1 and were designed with the aid of MAJIQ’s voila visualization tool.

### Immunoblot analyses

Immunoblot analysis was performed as described in Burke et al., 2019. Rabbit anti-GAPDH (Cell Signaling Technology: 2118L) was used at 1:2000. Anti-rabbit immunoglobulin G (IgG), horseradish peroxidase (HRP)–linked antibody (Cell Signaling Technology: 7074S) was used at 1:3000. Anti-mouse IgG, HRP-linked antibody (Cell Signaling Technology: 7076S) was used at 1:10,000. Histone H3 antibody (Thermo Fisher Scientific; NB500-171) was used at 1:1000. Crude nuclear and cytoplasmic fractionation was performed as described in Burke et al., 2017.

### Immunofluorescence and smFISH

smFISH was performed as described in Burke et al., 2019 and following the manufacturer’s protocol (https://biosearchassets.blob.core.windows.net/assets/bti_custom_stellaris_immunofluorescence_seq_protocol.pdf). GAPDH smFISH probes labeled with Quasar 570 dye (SMF-2026-1) were purchased from Stellaris. Custom IFNB1, IFNL1, SARS-CoV-2, and DENV2 smFISH probes were designed using Stellaris smFISH probe designer (Biosearch Technologies) available online at http://biosearchtech.com/stellaris-designer. Reverse complement DNA oligos were purchased from IDT (Data file S1). The probes were labeled with ATTO-633 using ddUTP-Atto633 (Axxora: JBS-NU-1619-633), ATTO-550 using 5-Propargylamino-ddUTP (Axxora; JBS-NU-1619-550), or ATTO-488 using 5-Propargylamino-ddUTP (Axxora; JBS-NU-1619-488) with terminal deoxynucleotidyl transferase (Thermo Fisher Scientific: EP0161) as described in Gaspar et al., 2017.

### Microscopy and image analysis

Coverslips were mounted on slides with VECTASHIELD Antifade Mounting Medium containing 4’,6-diamidino-2-phenylindole (DAPI) (Vector Laboratories; H-1200). Images were obtained using a wide-field DeltaVision Elite microscope with a 100× objective using a PCO Edge sCMOS camera. Between 10 and 15 Z planes at 0.2 μm per section were taken for each image. Deconvoluted images were processed using ImageJ with FIJI plugin. Z planes were stacked, and minimum and maximum display values were set in ImageJ for each channel to properly view fluorescence. Fluorescence intensity was measured in ImageJ. Single cells were outlined by determining the cell boundaries via background fluorescence and mean intensity was measured in the relevant channels. Imaris Image Analysis Software (Bitplane) (University of Colorado Boulder, BioFrontiers Advanced Light Microscopy Core) was used to quantify smFISH foci in nucleus and cytoplasm. Single cells were isolated for analysis by defining their borders via background fluorescence. Total foci above background threshold intensity were counted. Afterward, the nucleus marked with DAPI was masked, and foci were counted in the cell at the same intensity threshold cutoff, yielding the cytoplasmic foci count, from which the nuclear foci number could be determined.

### Figure generation

The model figure was created with BioRender.com.

