## Supplemental Figures and Legends for "RNase L-mediated RNA decay alters 3’ end formation and splicing of host mRNAs" (from another version of this preprint: version 1, 2022-01-28)

**Fig S1.**

(A) Distribution of intron/exon ratios in WT and RL-KO cells following mock or poly(I:C) lipofection of RNAs that are unchanged (padj > 0.5) in WT cells following poly(I:C) lipofection. B) Read coverage mapping to an unchanged RNA – TTC32. (C) RNAs that are downregulated (padj < 0.5, Log2Fold Change < -1). D) Read coverage mapping to a down-regulated RNA – TMEM60. (E) RNAs that are upregulated (padj < 0.5, Log2Fold Change > 1). F) Read coverage mapping to an up-regulated RNA – NOCT. Blue and red traces correspond to mapped reads on positive and negative strand respectively.

**Fig S2.**

(A) Distribution of DoG/exon ratios in WT and RL-KO cells following mock or poly(I:C) lipofection of RNAs that are unchanged (padj > 0.5) in WT cells following poly(I:C) lipofection. B) Read coverage mapping to an unchanged RNA – TTC32. (C) RNAs that are downregulated (padj < 0.5, Log2Fold Change < -1). D) Read coverage mapping to a down-regulated RNA – TMEM60. IGV trace scale was decreased 2.5x after autoscaling. (E) RNAs that are upregulated (padj < 0.5, Log2Fold Change > 1). F) Read coverage mapping to an up-regulated RNA – NOCT. Blue and red traces correspond to mapped reads on positive and negative strand respectively.

**Fig S3.**

(A) IGV tracks for NORAD in WT and RL-KO cells. Below, smFISH probes targeting the NORAD RNA or DoG are shown. (B) smFISH for NORAD CDS and DoG RNA in WT or RL-KO cells twelve hours following mock transfection or transfection with poly(I:C).

**Fig S4.**

(A) smFISH for IFNB1-CDS and IFNB1-DoG sixteen hours post-lipofection of poly(I:C), as shown in Fig 4C. (B) quantification of IFNB1 CDS in WT and RL-KO cells as represented in (A). (C-E) similar to (A) and (B) except analyzing IFNL1 CDS, IFNL1 Dog-1, and IFNL1 intron.

**Fig S5.**

(A) Co-smFISH for IFNB1 CDS and DOG-2 region in A549 WT and RL-KO cells sixteen hours post-lipofection with poly(I:C). (B) similar to (A) but for IFNL1 CDS and DoG-2 region.

**Fig. S6.**

(A) smFISH for IFNB1-CDS and IFNB1-DoG eight hours post-lipofection of poly(I:C). Red arrows indicate cells that contain high levels of both nuclear-retained IFNB1-CDS and IFNB1-DoG foci that mostly co-localize. White arrows indicate cells that do not contain high levels of IFNB1-DoG foci, which remain largely localized the IFNB1 genomic loci, and IFNB1-CDS is primarily localized to the cytoplasm. Yellow arrows demarcate cells that contain high levels of nuclear-localized IFNB1-CDS but not IFNB1-DoG foci. (B) Box plots displaying of the number of IFNB1-DoG foci localized to the nucleus or cytoplasm in WT or RL-KO cells as represented in (A). Scatter-plot of the ratio (nucleus/cytoplasm) of IFNB1-CDS foci (x-axis) and nuclear IFNB1-DoG foci in WT and RL-KO cells as represented in (A). (D) Scatter plots of the quantity of nuclear IFNB1-DoG foci (y-axis) and the quantity of nuclear IFNB1-CDS foci (x-axis) in WT and RL-KO cells as represented in (A). (E) smFISH for IFNL1-CDS and IFNL1-DoG in WT or RL-KO eight hours post-poly(I:C) lipofection.

**Fig S7.**

(A) smFISH for GAPDH and DENV2 mRNAs in WT A549 cells 48 hours post-infection with DENV2 (MOI=0.1). (B) similar to (A) but staining for PABP via IF.

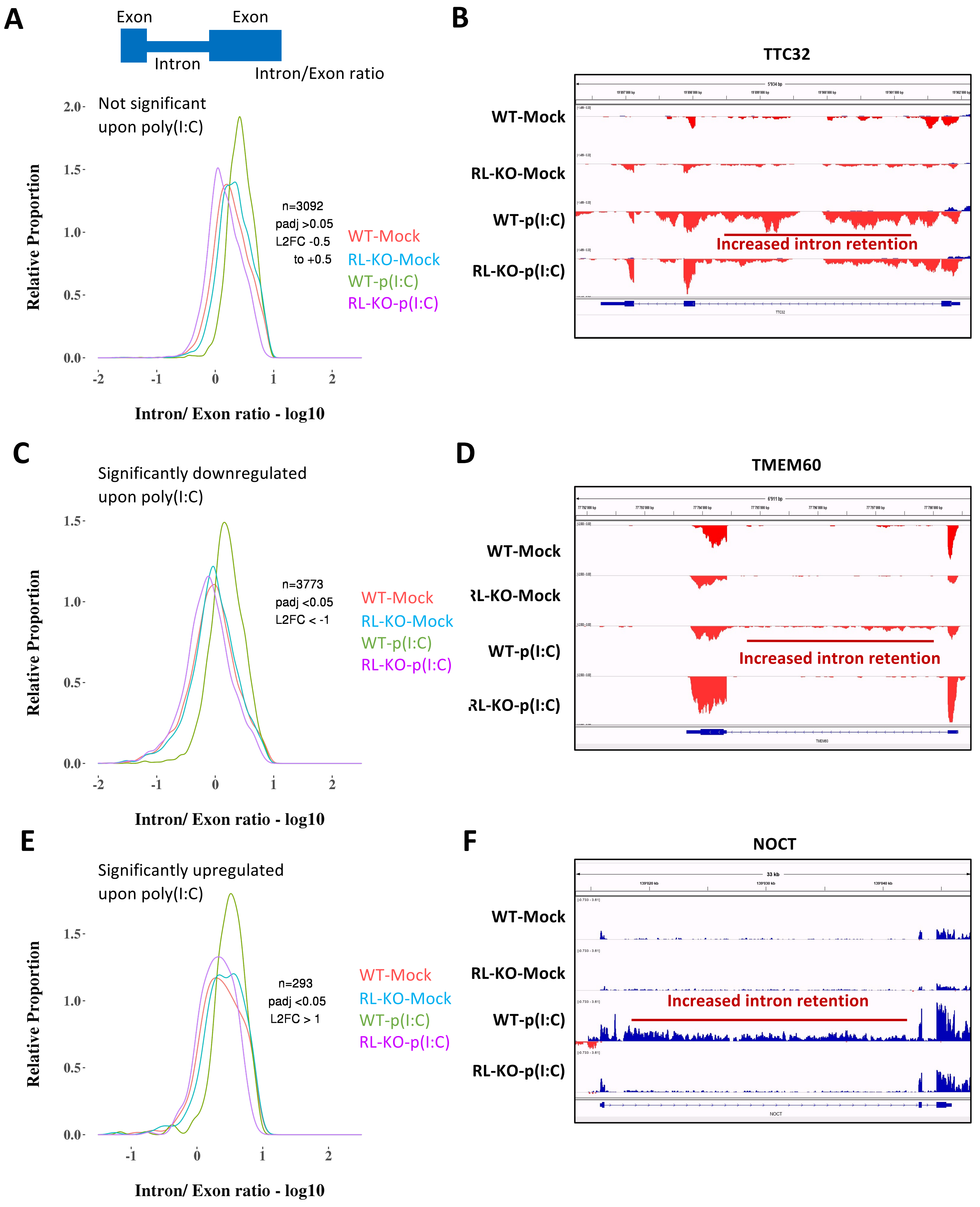
A
Exon
Exon
Intron
Intron/Exon ratio
Not significant
upon poly(I:C)
n=3092
padj >0.05
L2FC -0.5
to +0.5
WT-Mock
RL-KO-Mock
WT-p(I:C)
RL-KO-p(I:C)
Relative Proportion
Intron/ Exon ratio - log10
B
TTC32
WT-Mock
RL-KO-Mock
WT-p(I:C)
RL-KO-p(I:C)
Increased intron retention
C
Significantly downregulated
upon poly(I:C)
n=3773
padj <0.05
L2FC < -1
WT-Mock
RL-KO-Mock
WT-p(I:C)
RL-KO-p(I:C)
Relative Proportion
Intron/ Exon ratio - log10
D
TMEM60
WT-Mock
RL-KO-Mock
WT-p(I:C)
RL-KO-p(I:C)
Increased intron retention
E
Significantly upregulated
upon poly(I:C)
n=293
padj <0.05
L2FC > 1
WT-Mock
RL-KO-Mock
WT-p(I:C)
RL-KO-p(I:C)
Relative Proportion
Intron/ Exon ratio - log10
F
NOCT
33 kb
WT-Mock
RL-KO-Mock
WT-p(I:C)
RL-KO-p(I:C)
Increased intron retention

Figure S1

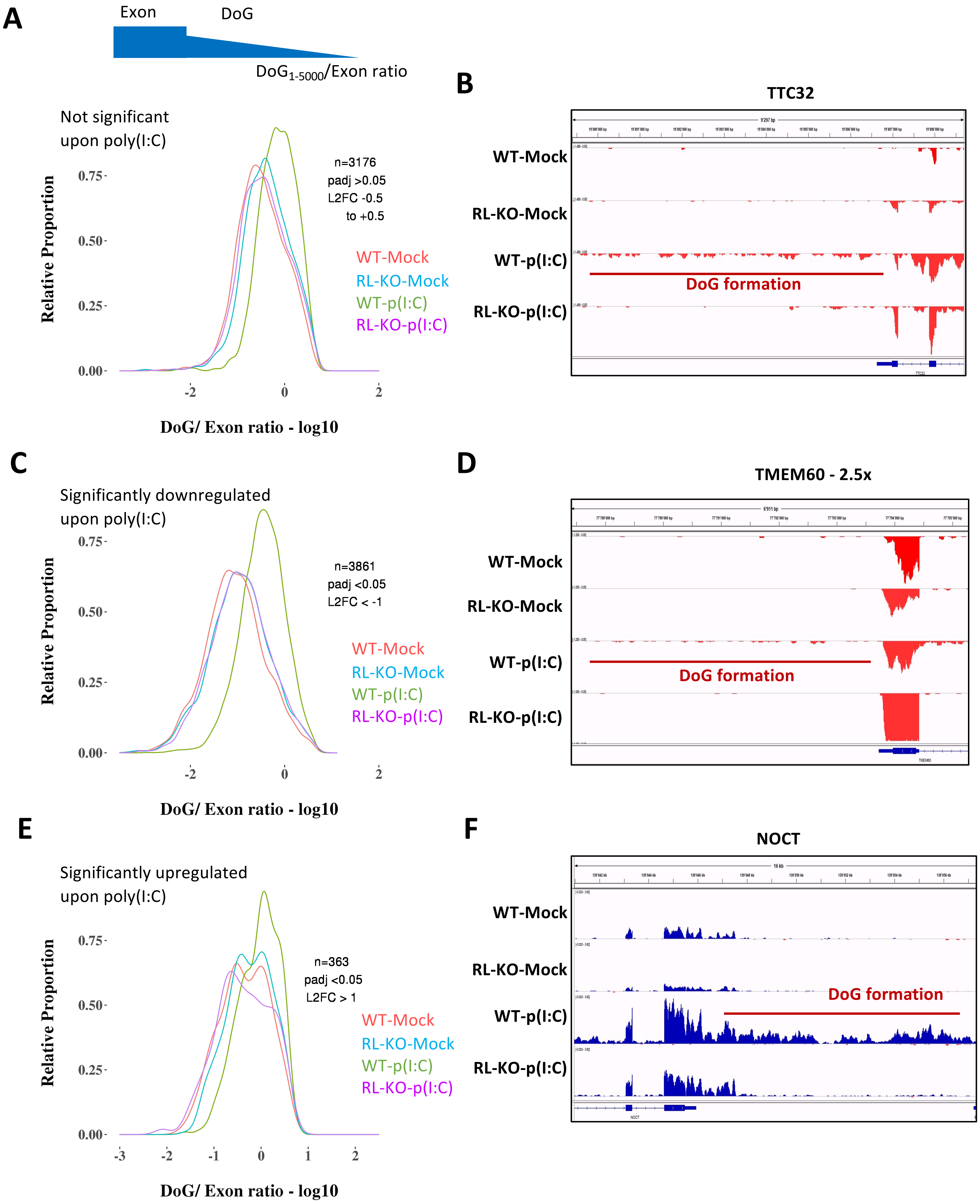
A
Exon
DoG
DoG1-5000/Exon ratio
Not significant
upon poly(I:C)
n=3176
padj >0.05
L2FC -0.5
to +0.5
WT-Mock
RL-KO-Mock
WT-p(I:C)
RL-KO-p(I:C)
Relative Proportion
0.75
0.50
0.25
0.00
-2
0
2
DoG/ Exon ratio - log10
B
TTC32
WT-Mock
RL-KO-Mock
WT-p(I:C)
DoG formation
RL-KO-p(I:C)
C
Significantly downregulated
upon poly(I:C)
n=3861
padj <0.05
L2FC < -1
WT-Mock
RL-KO-Mock
WT-p(I:C)
RL-KO-p(I:C)
Relative Proportion
0.75
0.50
0.25
0.00
-2
0
2
DoG/ Exon ratio - log10
D
TMEM60 - 2.5x
WT-Mock
RL-KO-Mock
WT-p(I:C)
DoG formation
RL-KO-p(I:C)
E
Significantly upregulated
upon poly(I:C)
n=363
padj <0.05
L2FC > 1
WT-Mock
RL-KO-Mock
WT-p(I:C)
RL-KO-p(I:C)
Relative Proportion
0.75
0.50
0.25
0.00
-3
-2
-1
0
1
2
DoG/ Exon ratio - log10
F
NOCT
WT-Mock
RL-KO-Mock
DoG formation
WT-p(I:C)
RL-KO-p(I:C)

Figure S2

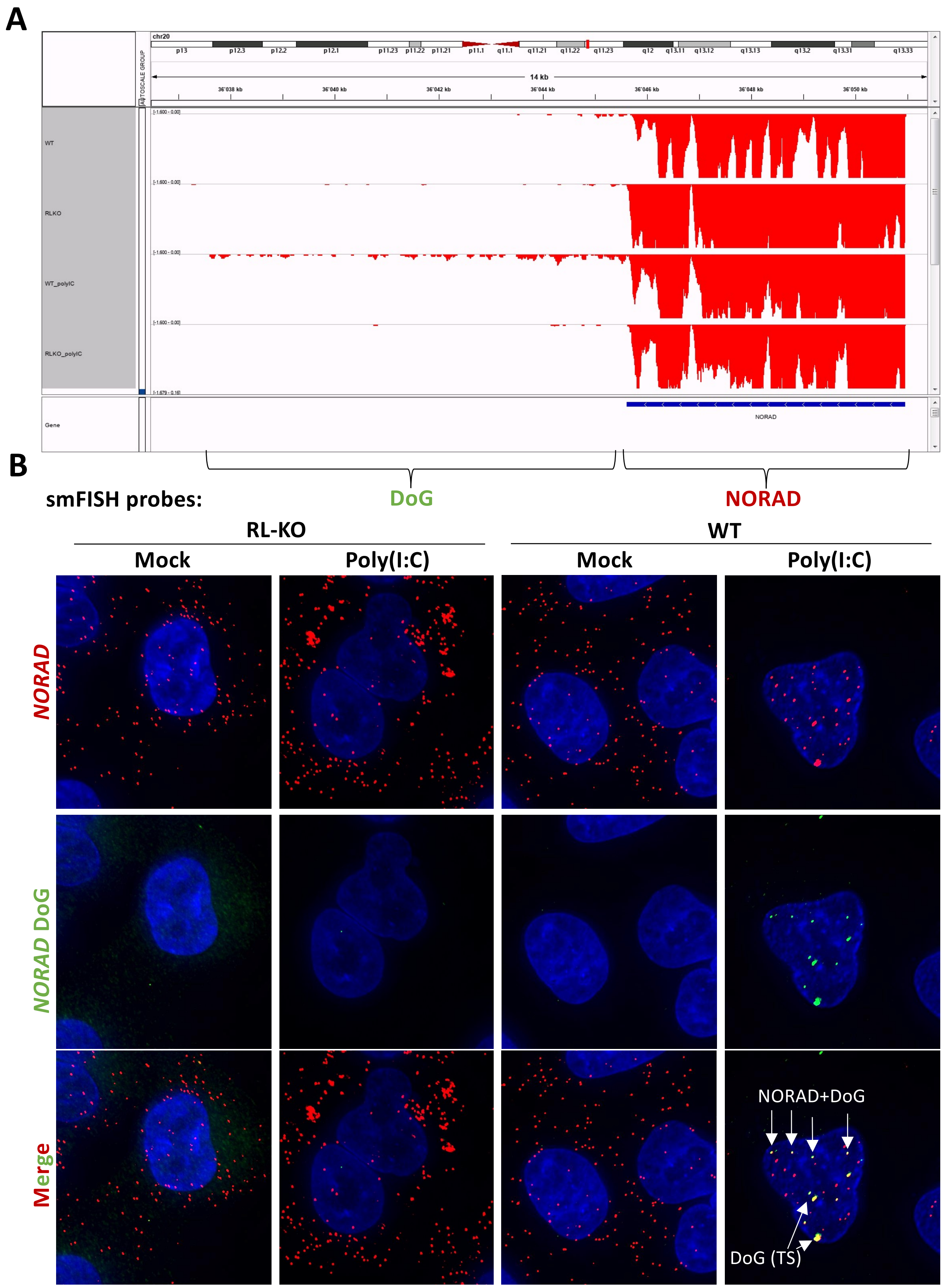
A
chr20
14 kb
WT
RLKO
WT_polyIC
RLKO_polyIC
Gene
NORAD
B
smFISH probes:
DoG
NORAD
RL-KO
WT
Mock
Poly(I:C)
Mock
Poly(I:C)
NORAD
NORAD DoG
Merge
NORAD+DoG
DoG (TS)

Figure S3

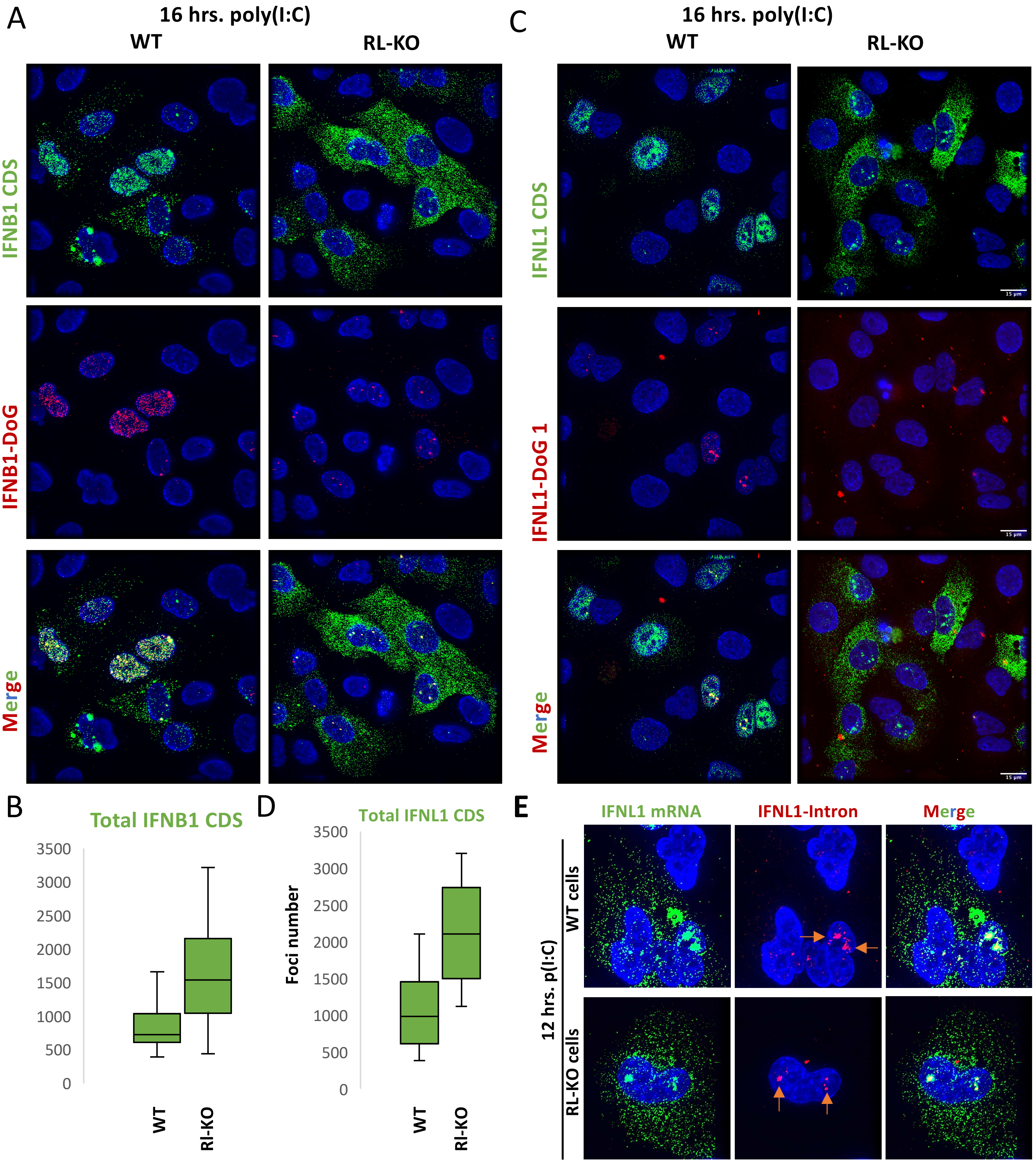
A
16 hrs. poly(I:C)
WT
RL-KO
IFNB1 CDS
IFNB1-DoG
Merge
C
16 hrs. poly(I:C)
WT
RL-KO
IFNL1 CDS
IFNL1-DoG 1
Merge
B
Total IFNB1 CDS
3500
3000
2500
2000
1500
1000
500
0
WT
RI-KO
D
Total IFNL1 CDS
Foci number
3500
3000
2500
2000
1500
1000
500
0
WT
RI-KO
E
IFNL1 mRNA
IFNL1-Intron
Merge
12 hrs. p(I:C)
WT cells
RL-KO cells

Figure S4

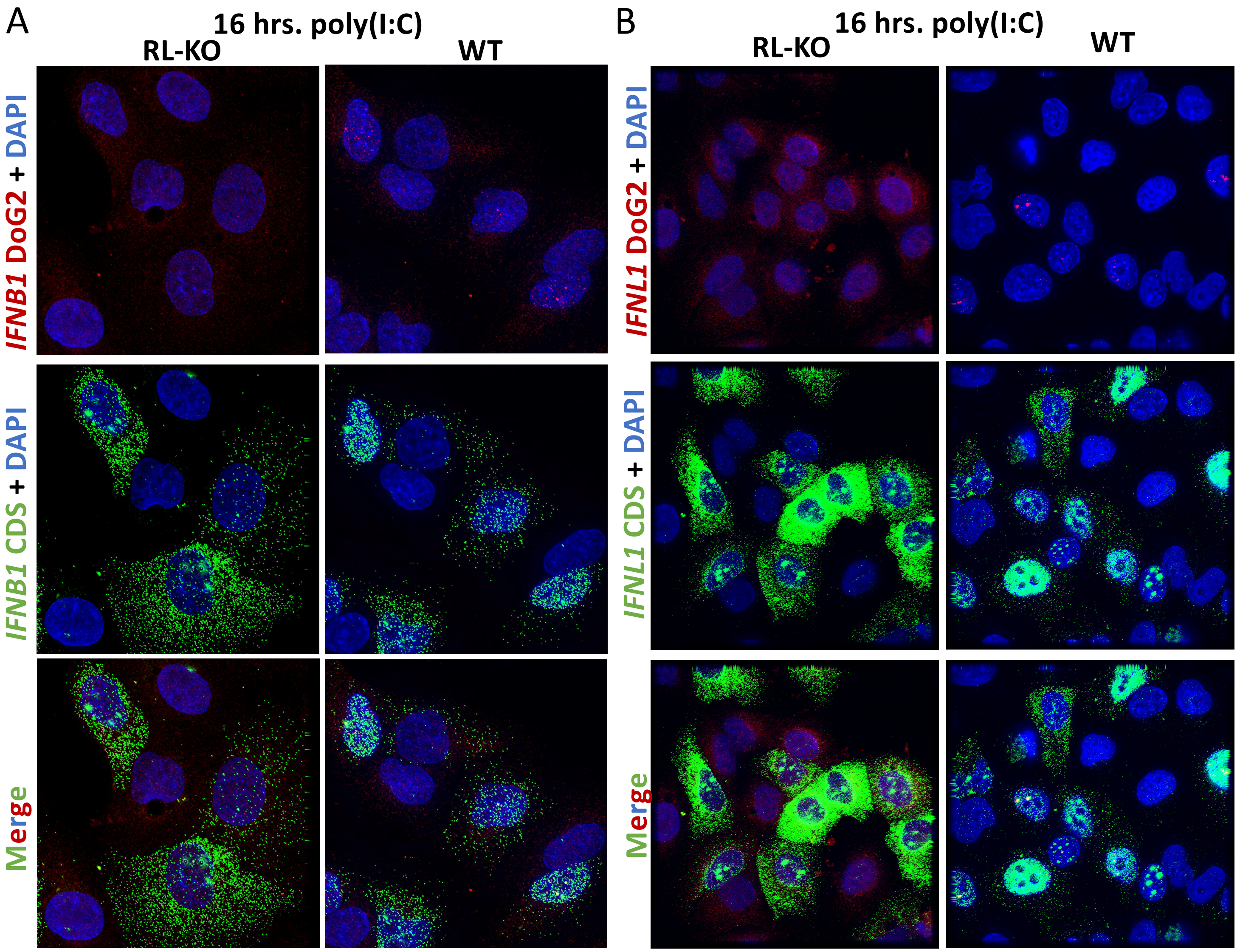
A
16 hrs. poly(I:C)
RL-KO
WT
IFNB1 DoG2 + DAPI
IFNB1 CDS + DAPI
Merge
B
16 hrs. poly(I:C)
RL-KO
WT
IFNL1 DoG2 + DAPI
IFNL1 CDS + DAPI
Merge

Figure S5

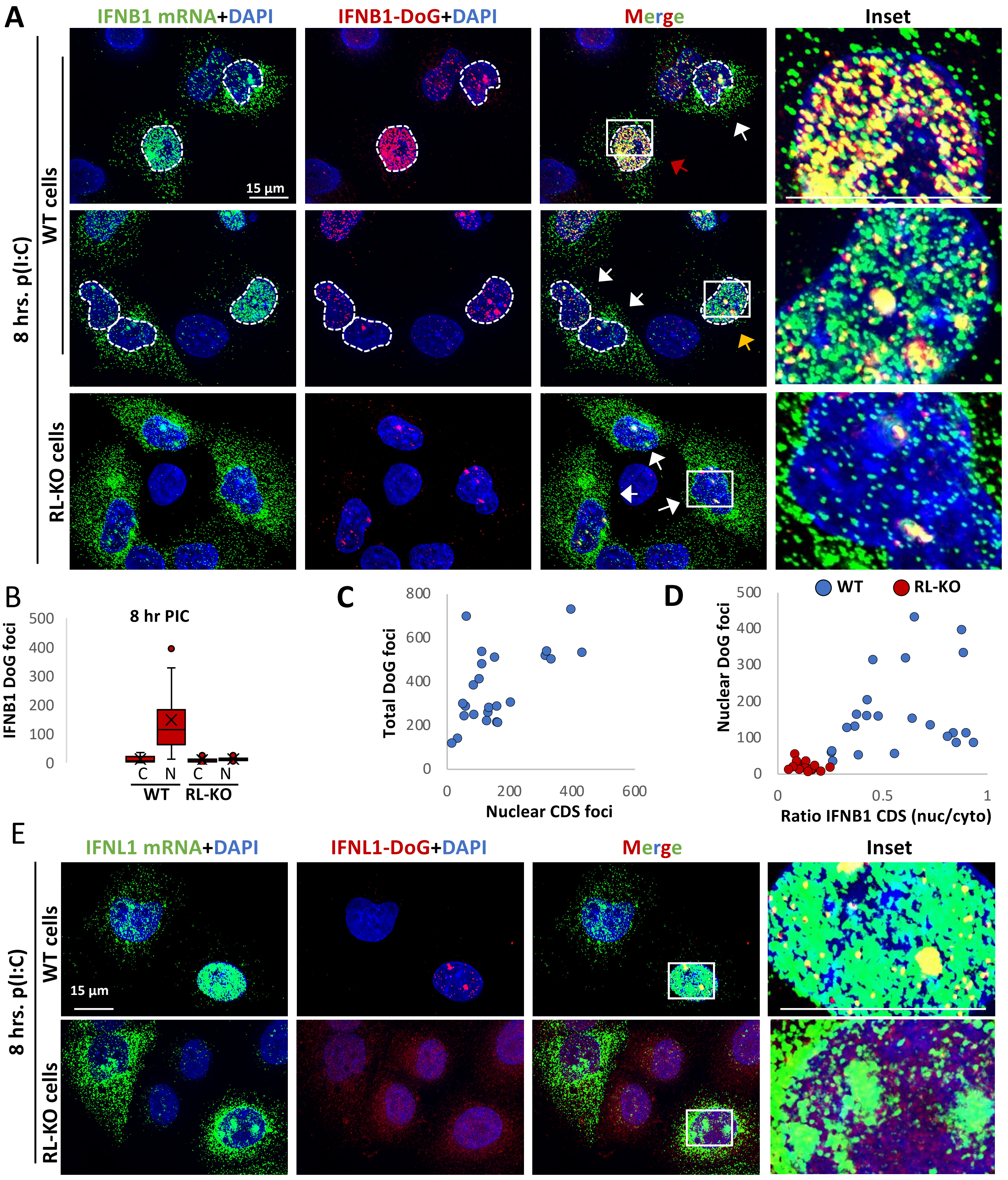
A
IFNB1 mRNA+DAPI
IFNB1-DoG+DAPI
Merge
Inset
8 hrs. p(I:C)
WT cells
RL-KO cells
15 µm
B
8 hr PIC
IFNB1 DoG foci
500
400
300
200
100
0
C
N
C
N
WT
RL-KO
C
Total DoG foci
800
600
400
200
0
0
200
400
600
Nuclear CDS foci
D
WT
RL-KO
Nuclear DoG foci
500
400
300
200
100
0
0
0.5
1
Ratio IFNB1 CDS (nuc/cyto)
E
IFNL1 mRNA+DAPI
IFNL1-DoG+DAPI
Merge
Inset
8 hrs. p(I:C)
WT cells
RL-KO cells
15 µm

Figure S6

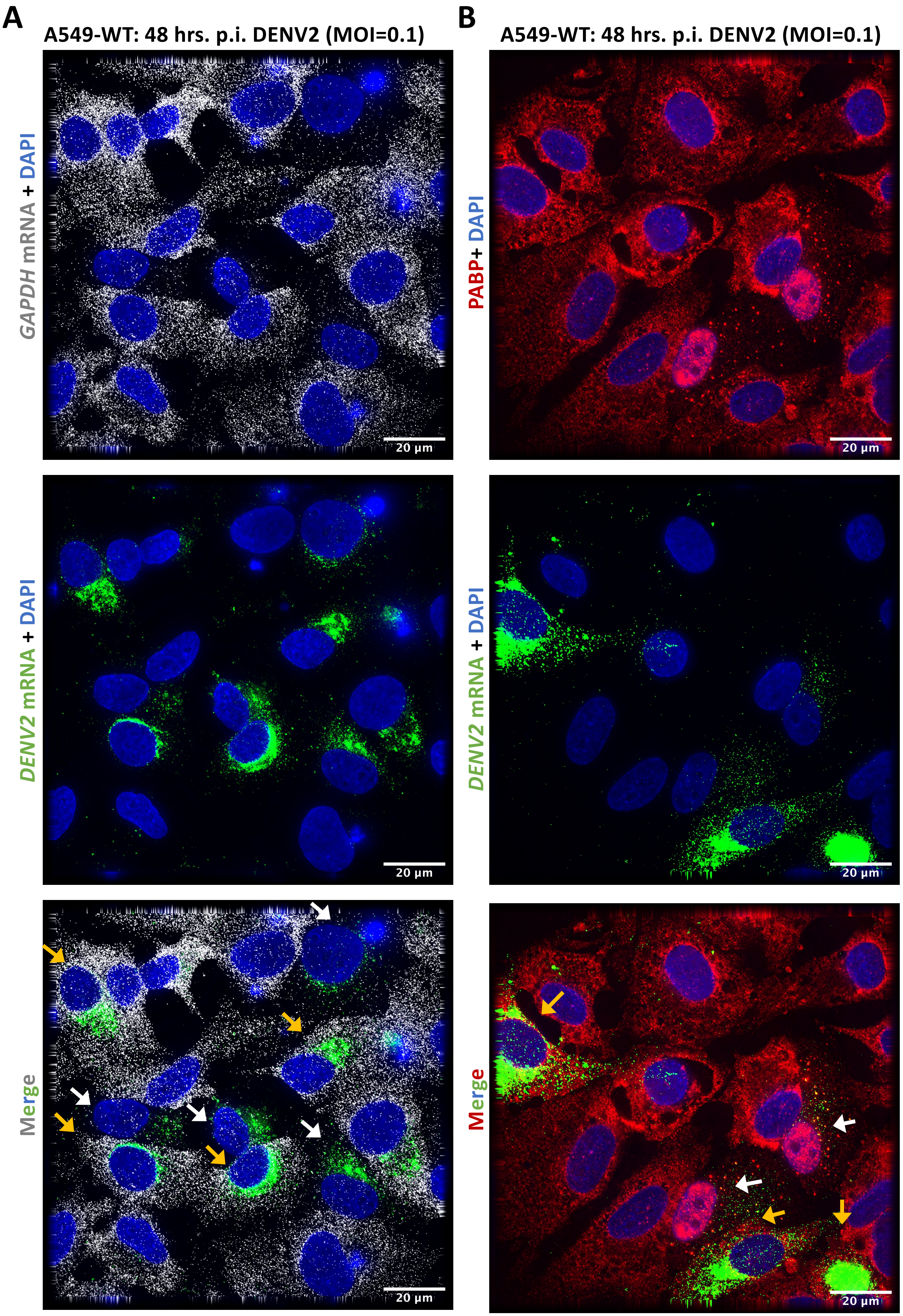
A
A549-WT: 48 hrs. p.i. DENV2 (MOI=0.1)
GAPDH mRNA + DAPI
DENV2 mRNA + DAPI
Merge
20 µm
B
A549-WT: 48 hrs. p.i. DENV2 (MOI=0.1)
PABP+ DAPI
DENV2 mRNA + DAPI
Merge

Figure S7
